# Pathfinder: protein folding pathway prediction based on conformational sampling

**DOI:** 10.1101/2023.04.20.537604

**Authors:** Zhaohong Huang, Xinyue Cui, Yuhao Xia, Kailong Zhao, Guijun Zhang

**Affiliations:** College of Information Engineering, Zhejiang University of Technology. His research interests include bioinformatics, intelligent information processing, and optimization theory; College of Information Engineering, Zhejiang University of Technology. Her research interests include bioinformatics, intelligent information processing, optimization theory; College of Information Engineering, Zhejiang University of Technology. His research interests include bioinformatics, intelligent information processing, optimization theory

**Keywords:** Protein folding, Intermediate state, Conformational sampling, Pathway prediction

## Abstract

The study of protein folding mechanism is a challenge in molecular biology, which is of great significance for revealing the movement rules of biological macromolecules, understanding the pathogenic mechanism of folding diseases, and designing protein engineering materials. Based on the hypothesis that the conformational sampling trajectory contain the information of folding pathway, we propose a protein folding pathway prediction algorithm named Pathfinder. Firstly, Pathfinder performs large-scale sampling of the conformational space and clusters the decoys obtained in the sampling. The heterogeneous conformations obtained by clustering are named seed states. Then, a resampling algorithm that is not constrained by the local energy basin is designed to obtain the transition probabilities of seed states. Finally, protein folding pathways are inferred from the maximum transition probabilities of seed states. The proposed Pathfinder is tested on our developed test set (34 proteins). For 5 widely studied proteins, we correctly predicted their folding pathways. For 25 partial biological experiments proteins, we predicted folding pathways could be further verified. For the other 4 proteins without biological experiment results, potential folding pathways were predicted to provide new insights into protein folding mechanism. The results reveal that structural analogs may have different folding pathways to express different biological functions, homologous proteins may contain common folding pathways, and α-helices may be more prone to early protein folding than β-strands.

## 1 Introduction

The splendid computational success of AlphaFold2 [1] and RoseTTAFold [2] in protein structure prediction may have solved the static single domain protein folding problem [3]. The AlphaFold database [4] and recent predictions of more than 200 million protein structures provide reference structure information for nearly every known protein [5]. Although, the greatly improved prediction of protein 3D structure from sequence achieved by the AlphaFold2 has already had a significant impact on biological research [6], but challenges remain[7, 8]. Almost all computational methods are unable to predict accurate protein folding pathway [9-11]. This is because protein folding is a dynamic process of exploring the overall energy landscape and locating heterogeneous local energy basins to obtain its functional structure and conformations [12]. The study of folding mechanism is of great significance for the formation of inclusion bodies [13] and for revealing the second genetic code [14]. Many pathological conditions are also fundamentally rooted in the misfolding, aggregation, and accumulation that occurs in protein folding [15], such as Alzheimer’s disease [16], Parkinson’s disease [17], and other diseases. Understanding the folding mechanism can provide important implications for the treatment of these diseases [18], as well as facilitate the design of proteins with unique functional characteristics [19, 20], and the exploration of protein allosteric [21]. The conformational heterogeneity of different states in protein folding, such as unfolded state [22], misfolded state [23], intermediate state [24], and transition state [25], is crucial for an accurate understanding of folding mechanisms [26].

There are biological experimental methods to explore protein intermediate states and folding pathways[27-32], such as hydrogen deuterium exchange mass spectrometry [33] and circular dichroism spectrum [34]. However, biological experimental methods are difficult to obtain high-resolution spatial and temporal data on the folding process. This is because the biological process by which proteins fold into their unique native state occurs within seconds to minutes [35], and metastable conformations are more difficult to detect due to their short lifetime and low occupancy [36]. This makes it challenging to explore intermediate states with biological experimental methods.

Computational simulation of protein folding can make up for the deficiency of biological experimental methods, and is an effective way of studying protein folding pathways [37]. Molecular Dynamics Simulation (MD) is one of the methods for computationally simulating protein folding, which can simulate the complete folding process of small molecules [38]. MD combined with Markov models can analyze folding and functional dynamics in long trajectories [39]. Machine learning facilitates protein folding simulations by extracting essential information and sampling of rare events from large simulated datasets [40]. Convolutional neural networks learn continuous conformational representations generated from protein folding simulations to predict biologically relevant transition paths [41]. Integrating biological experimental structural constraints into MD models significantly explored protein dynamics trajectories [42]. The current mainstream method is combining biological experiments and MD to explore protein folding mechanism. However, MD are more applied to the simulation of short trajectories between states, and it is still challenging to simulate the complete folding process [37]. Our recent work (PAthreader) [43] identifies remote homologous structures based on the three-track alignment of distance profiles and structure profiles originated from Protein Data Bank (PDB) [44] and AlphaFold database by deep learning. Based on the recognized homologous templates, PAthreader further explored protein folding pathways by identifying folding intermediates, but it has limitations on proteins that lack remote homologous template information.

Conformational sampling algorithms such as Monte Carlo (MC) can be applied to folding simulations of template-free and larger proteins [45]. The CA-CB side chain model combined with MC kinetics to identify the protein folding pathway and the interaction pairs during the folding process [46, 47]. Conformational sampling tends to fall into local energy traps. Recently, we have developed several methods (MMpred [48], SNfold [49]) to compensate for this deficiency. The MMpred aims to explore the complete energy landscape and improve sampling efficiency, which can be beneficially applied to explore heterogeneous conformations. MMpred has localized promising energy basins in parallel on multiple trajectories, combining a greedy search strategy with distance-constrained information to infer the final structure. SNfold overcomes high-energy barriers and avoids resampling of exploration regions to obtain diverse heterogeneous conformations in the energy landscape. These state-of-the-art conformational sampling algorithms are mainly used in protein structure prediction. However, the idea of exploring multiple states can be effectively applied to protein folding pathway prediction.

In this work, we propose a protein folding pathway prediction algorithm (Pathfinder) based on conformational sampling. We obtain the structural information of the seed states through large-scale sampling and explore state transition probabilities through resampling. Pathfinder captures the information (seed states, sampling states and transition probabilities) to predict folding pathways. Pathfinder is tested on our developed test set (34 proteins). Our predicted folding pathway were consistent with biological experiments for 5 widely studied proteins including the B1 domain of protein L and protein G, two SRC homology 3 domains, and the LysM domain. Because of the imperfection of existing protein folding data. For 25 partial biological experiments proteins, we predicted folding pathways could be further verified. For the other 4 proteins without biological experiment results, potential folding pathways were predicted to provide new insights into the protein folding mechanism. Analyzing of the above results, we found some protein folding mechanisms.

## 2 Methods

Protein conformational sampling can provide new ways to explore folding pathways. In this study, we hypothesized that the protein folding information from unfolded state to folded state may be implied in the conformational sampling process in the energy landscape [46, 50], and that the maximum probability path of state transition corresponds to the folding pathway. Based on the above assumptions and inspired by hidden Markov model, we predict protein folding pathways by the transition probability between metastable states inferred from sampled conformations. Here, the metastable states located in local energy basins are named as ‘Seed states’ (cyan structures, Figure 1), where the states in shallow basin of folding pathway are called ‘Intermediate states’.

**Figure 1.**
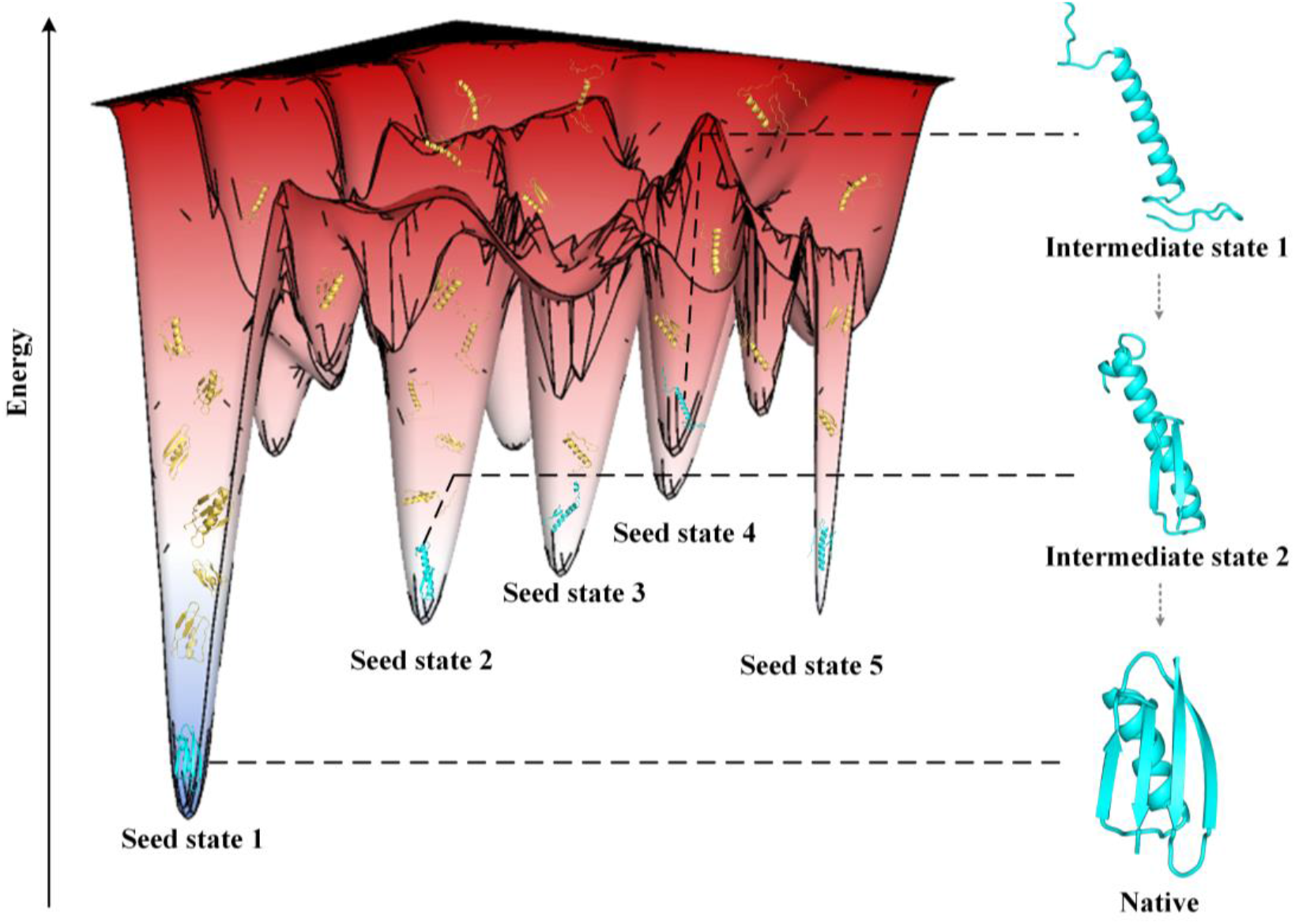
Schematic diagram of folding pathway prediction based on conformational sampling. The yellow structures are sampling states and cyan structures are seed states. The dotted arrows indicate the implicit transition between intermediate states. Sampling states are obtained by the large-scale conformational sampling. The seed states and their transition tendency are inferred by the resampling algorithm.

### 2.1 Overview

The pipeline of Pathfinder is shown in Figure 2, consists three stages: (A) seed generation, (B) transition probability exploration and (C) folding pathway inference. The input is the query sequence and native structure from PDB (or predicted model by AlphaFold2 if there is no crystal structure in PDB). The output is the predicted protein folding pathway. Guided by the energy function of *ClassicAbinitio* protocol in Rosetta [51, 52], fragment assembly-based Metropolis Monte Carlo (MMC) algorithm [53] is used for conformational sampling in the stages (A) and (B). The fragment library is built by the Robetta fragment server (http://old.robetta.org/). In the stage (A), a large-scale conformational sampling algorithm is used to obtain a mass of decoys. The cluster centroid obtained by clustering is selected as the seed state. In the stage (B), based on a modified energy function, conformational resampling is not constrained by local energy traps to explore transition probabilities between seed states. Finally, the protein folding pathways are inferred by transition probabilities using a dynamic programming algorithm in the stage (C).

**Figure 2.**
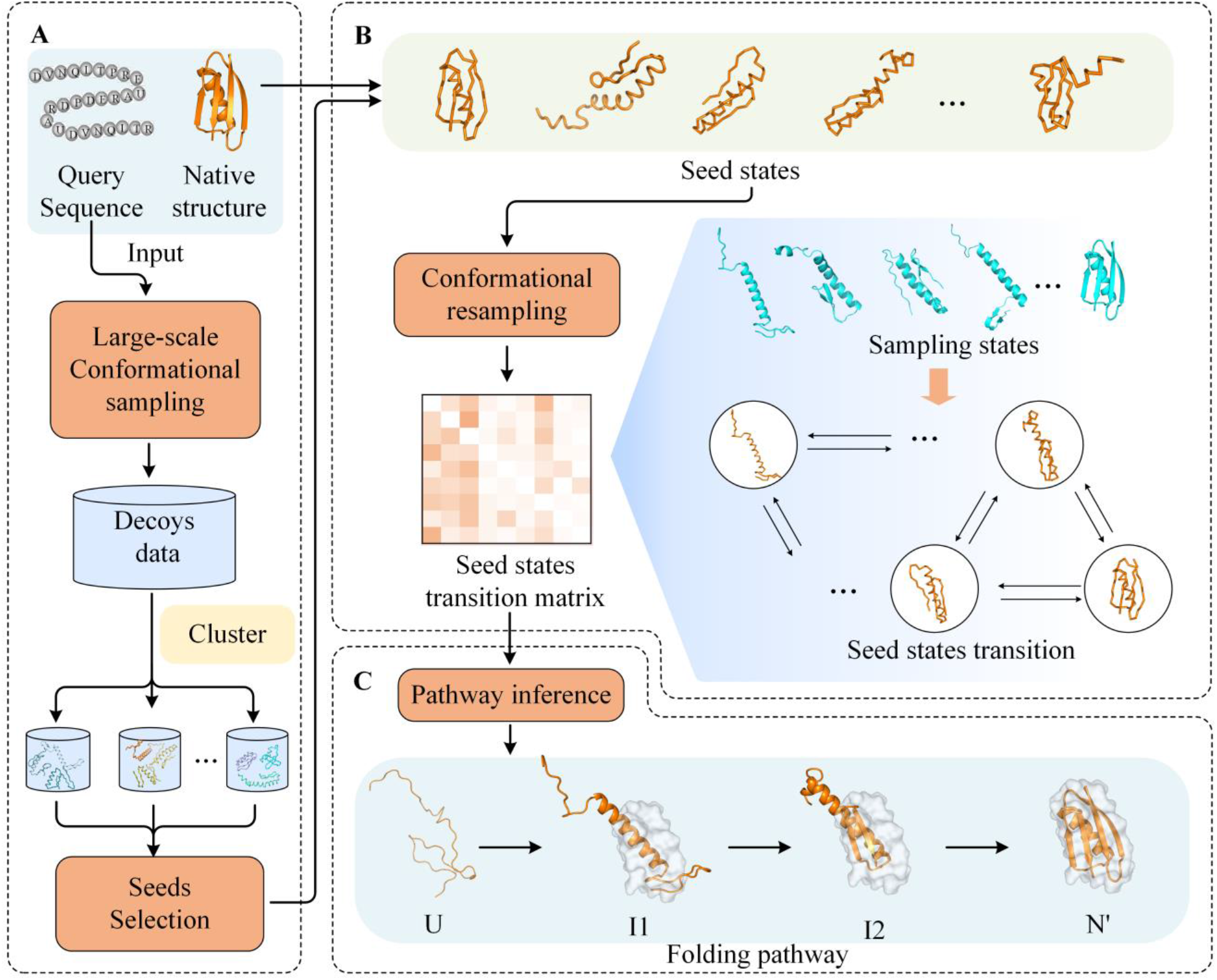
The pipeline of Pathfinder. (A) Seed generation. Sampling of large-scale conformational space by input sequences. Cluster and output the seed states. (B) Transition probability exploration. The seed states in this stage consists of the seeds obtained in stage A and the input native structure. (C) Folding pathway inference. The folding pathway starts from the unfolded state and passes through several intermediate states to a near-native state (N’), which is the closest conformation to native during sampling.

### 2.2 Seed generation

We use a large-scale conformational sampling with *G* MMC trajectories to explore energy basins with high repetition rates in folding pathways. Each MMC trajectory generates about 360,000 conformations, where accepted conformations are saved as decoys. There are at least 13000 decoys from all MMC trajectories for cluster. We cluster decoys into centroids using Spicker [54], and every 13,000 decoys are clustered into *S* centroids. Because of redundancy among centroids, we merge centroids with TM-score>*τ* to generate *N* seed states. In particular, the lowest global energy basin is represented in the seed state 1 (illustrated in Figure 1) of the input structure.

### 2.3 Transition probability exploration

#### 2.3.1 Modified energy function

Different from the large-scale sampling, the purpose of resampling algorithms is to explore the transition propensity of the seed states, rather than to simulate de novo protein folding. Because there are masses of energy barriers in the energy landscape, the conformational sampling has the defect that it is difficult to jump out after entering the local energy basin, which leads to low sampling efficiency. Therefore, we construct a modified energy function to facilitate sampling state transitions by raising the energy basin and lowering the energy barrier as illustrate in Figure 3.

**Figure 3.**
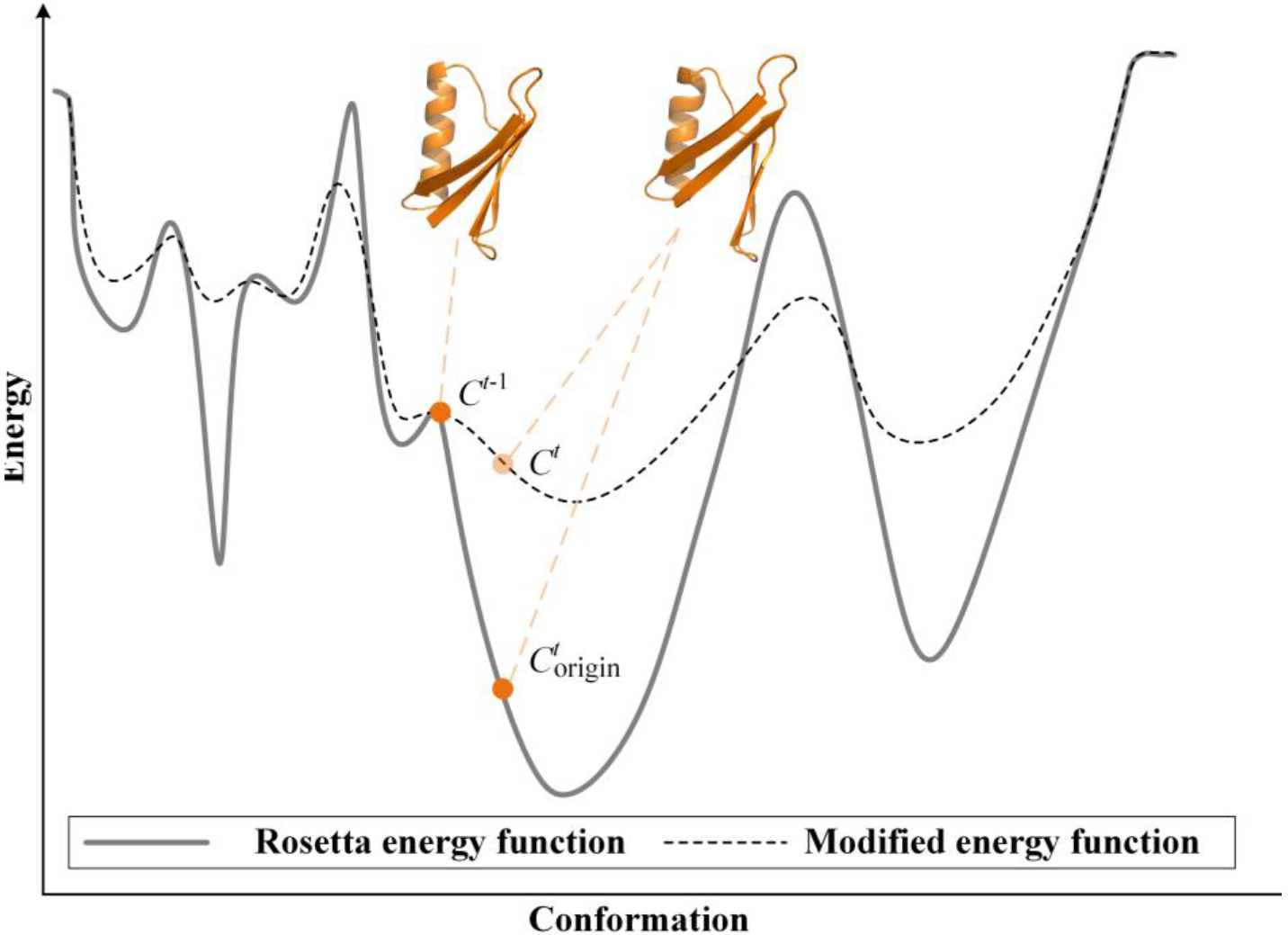
Schematic of the modified energy landscape. After modification, the energy basin is raised and the energy barrier is lowered. And the relatively smooth energy landscape makes transitions between states easy.

The *C^t^* is the *t*-th conformation accepted in modified energy landscape, the 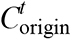 is the original conformation in the unmodified energy landscape, and *C^t-^*^1^ is (*t*-1)-th conformation. Based on the modified energy function, *C^t^* escapes the local basin more easily than 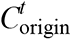 The modified energy function *f* (*C^t^*) guiding resampling is defined as:

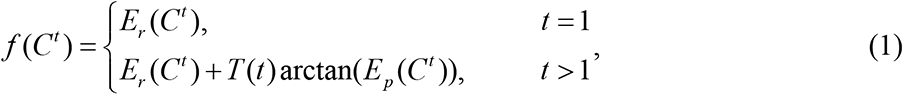

where *E_r_* (*C^t^*) is Rosetta energy function and *E_p_* (*C^t^*) is an energy function which designed to modify the original energy function. *T* (*t*) is reduced as the number of samples increases to offset the large energy gap between the unfolded state and the folded state.

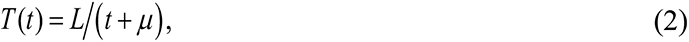

where *L* is the length of the protein, the *μ* is the initial value to avoid excessive energy at the beginning of sampling. The energy function *E_p_* (*C^t^*) is designed as:

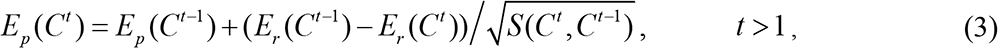

where *E_p_* (*C^t^*^−1^) is the previously accumulated energy function to maintain enough energy to rush out of the energy basin. *S* (*C^t^*, *C^t^*^−1^) is the dihedral angles difference between *C^t^* and *C^t^*^−1^, designed as:

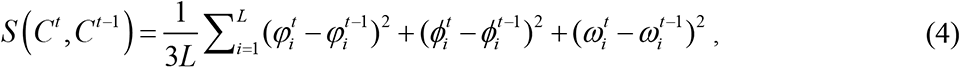

where 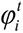, 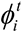 and 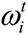 are the dihedral angles of *i* residue of *t*-th conformation. The modified energy function was compared with the benchmark conformational sampling algorithm (Supplementary Figure 1). The results show that the modified energy function can explore wider energy basins and guide the conformational sampling process more quickly.

#### 2.3.2 Seed states transition generation

In the funnel model, high-energy barriers exist around local energy basins, and random jumps in sampling points between basins contain transition information [35]. The resampling algorithm locates energy basins for sampling in conformational space based on sampling states and DMscore cutoff *η*. DMscore [48] is a structural similarity metric we previously developed, focusing on secondary structure to determine the extent of local energy basins. Based on the modified energy function, the resampling algorithm utilizes conformational sampling to obtain potential seed state transition probabilities. Inspired by hidden Markov models, the observation state is a representation of the hidden unknown state. The state maximum probability path can be obtained by constructing the model and the observation state. However, state transition path inference needs to obtain a continuous sequence of observation states and cannot be directly used in random image sampling methods. Therefore, we generate a mass of sampling states to obtain state transitions by comparing their structural similarity with the seed states. MMC trajectories are different from the continuous trajectories of molecular dynamics simulations, including random conformational structure transitions. We define the state transition frequency matrix as **B** = {*b_ij_*}, where *b_ij_* is obtain by resampling algorithm. We consider the sampled trajectory to enter this state region when the DMscore between the sampling conformation and seed *i* is greater than *ξ*. At the same time, we designed the unidirectionality of the transformation, that is, when the transmission from *i* to *j* is the reverse of the previous transmission, the transmission frequency is not calculated, which helps to reduce the background noise generated by random sampling. The transition frequency of the seed states is calculated by resampling. We further process the **B** to get the transition probability matrix **A** = {*a_ij_* }, where 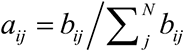

### 2.4 Folding pathway inference

Based on the seed states transition probability matrix **A**, protein folding pathways are inferred using a dynamic programming algorithm. The folding pathways are represented in the order of seed states, which are inferred from the transition probability matrix. Based on the *N* seed states obtained above, the optimal path is defined as 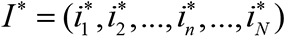, where 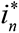 is the optimal path from 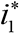 to 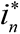. The 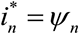 is defined as:

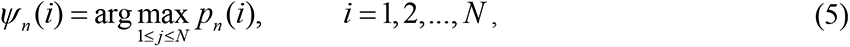

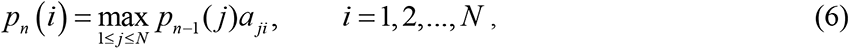

where *p_n_* (*i*) is the maximum transition probability to 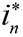, and *a _ji_* is the transition probability from seed *i* to seed *j* .Because of the imperfection of the Rosetta force field, it is difficult to sample protein conformations to the native state. Therefore, the known native structure is used as the end point of the folding (*ψ*_1_ (*i*) = 1), and the folding pathway *I* * is reversely inferred.

### 2.5 Evaluation metric

Related studies have shown that the logarithm of experimental protein folding rates depends on the local geometry and topology of the protein’s native state [55]. Contact order is a metric of protein topology complexity and stability, reflecting the relative importance of local and nonlocal contacts to protein structure [56]. Contact order has a statistically significant relationship with protein folding dynamics and was used in this work to assess folding dynamics in intermediate states [57], which is defined as:

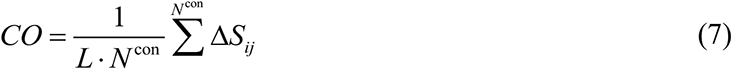

where *N*^con^ is the number of residues whose distance between them is less than 8Å. The *ΔS_ij_* is the sequence separation between residue *i* and *j*, and *L* is the length of the protein. On this basis, the residue contact order of *i*-th *R*_co_ (*i*) was designed to assess the local folding completion of intermediate states:

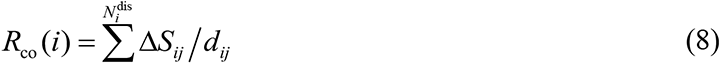

where 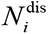 is the number of residues whose distance are less than 20Å from *i*-th residue. The *d_ij_* is the distances between residue *i* and *j.* The contact order can evaluate the degree of folding completion through the intermediate state structure information. The residue contact order can capture the folding nucleus information and key residue information during the folding process.

## 3. Result

### 3.1 Test set

The test set we collected includes 34 proteins. The folding data of 6 proteins are collected from the HDX experimental database of Start2Fold [58].19 proteins are collected from the Protein Fold Database (PFDB) [59], the other 5 proteins come from our collection of related protein folding research papers. The above 30 proteins are included with known protein folding data, and related research papers are in the appendix (Supplementary Figure2, 3 and 4). We also collected 4 proteins with no experimental folding information.

### 3.2 Predicted folding pathway on proteins with experimental validation

#### 3.2.1 The GB1 and LB1

The IgG-binding B1 domain of protein G (GB1, Figure 4a) and IgG-binding B1 domain of protein L (LB1, Figure 4d) are often used as model proteins for folding mechanism studies. GB1 has a wide range of biomedical uses and studies of protein folding and stability [60], and extensive experimental results and computational simulations exploring the complete folding pathway[46]. Both GB1 and LB1 contain an α-helix and four β-strands, but their sequence similarity is low and their folding pathways differ. Related studies have shown that the hairpin structure (folding nucleus) plays a crucial role in global folding [46].

**Figure 4.**
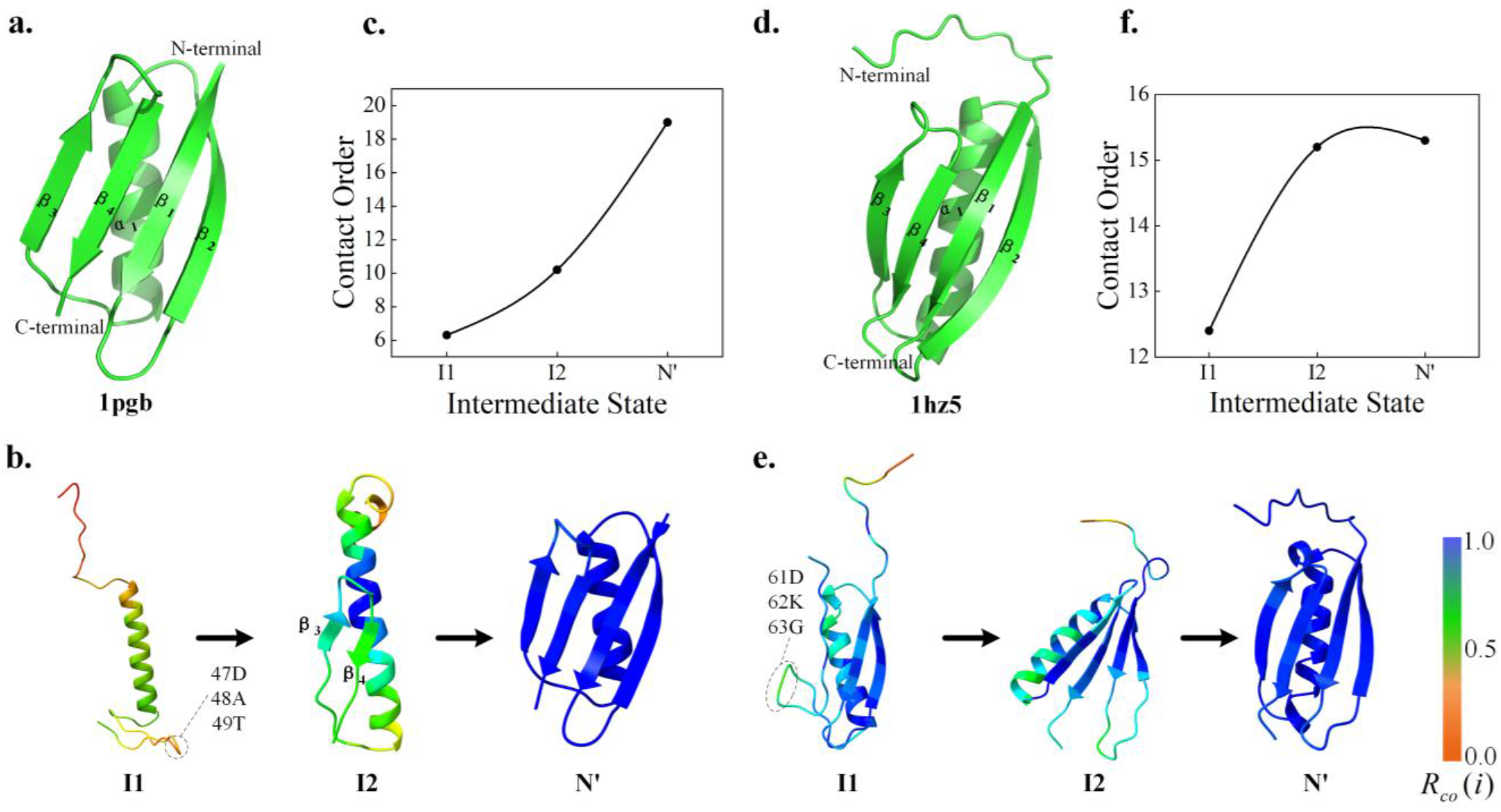
Folding pathway of GB1 (PDB ID:1pgb) and LB1 (PDB ID:1hz5). **(a)** and **(d)** are the native structure of GB1 and LB1. **(b)** and **(e)** are the folding pathway of GB1 and LB1 including intermediate states. **(c)** and **(f)** are the contact orders of the intermediate states. Residue contact order values are normalized and represented on the structure as color.

The predicted protein folding pathways are shown in Figure 4b and e. The result shows that GB1 first forms an α-helix, meanwhile the β-turns (47D, 48A, 49T) at the C-terminus has started to form, which may be a sign that I2 of GB1 (called G_I2_) start to form. Then the β_3_ and β_4_ formed as shown in G_I2_. In addition, the β_1_ and β_2_ of G_I2_ is represented in helical, which is different from the native structure in Figure 4a. This is because the unstable structure in the intermediate state usually exists as a disordered region, and it may be replaced by secondary structures such as helices and loops in fragment assembly. Finally, a hairpin structure composed of β_1_ and β_2_ is formed at the N-terminus (as shown in N’). Our GB1 predictions are complete consistent with biological experiments [61-63] and contain structural information for the folding stage. As shown in Supplementary Figure 5, the folding pathway of GB1 can be clearly observed through the residue contact orders in the intermediate state. Therefore, after normalizing the residue contact orders, the color of the scale in the lower right corner of Figure 4 shows the folding degree of the residues.

As shown in 4f, the I1 of LB1(called L_I1_) suggest that the β_1_ and β_2_ hairpin structures at the N-terminal are formed earlier than the β_3_ and β_4_ structures, which was different from the folding mechanism of GB1. After β_3_ and β_4_ folding, the loop of LB1 folds to stabilize. Similarly, C-terminal β turns (61D, 62K, 63G) and β_4_ start to form in the L_I1_. The predicted results of LB1 are almost consistent with the biological experiments [64], but the order of helix and N-terminal hairpin structure formation is missing. By analyzing the conformational structure of the sampling process (Supplement Figure 6), we found that the sampling occupancy rate of these two structures is low, and the clustering structure is mainly the process of β_3_ and β_4_ and loop. This is because the fragment assembly method completes both the helix and sheet folding early in conformational sampling.

As shown in Figure 4c and f, the contact order of their intermediate states showed an upward trend, reflecting the degree of globular protein folding. The folding of L_I2_ to N’ is mainly affected by loop adjustment, and the degree of folding is much lower than that of β-strand formation from the I1 of LB1 to L_I2_. In addition, related studies have shown that the formation order of the hairpin structure is crucial to the folding rate and stability [62]. Inspired by this difference in folding mechanism, the GB1 mutant NuG2 (PDB ID: 1mi0) was designed [64]. We also predicted the folding pathway of the NuG2 as shown in Supplementary Figure 2. The results showed that the two β-turns formed almost simultaneously at the beginning of folding. The above results suggest that structural analogs may have different folding pathways and provide material for protein design.

#### 3.2.2 SRC Homology 3 Domain

The study on the folding of SRC Homology 3 Domain provides extremely valuable information for the molecular mechanism of amyloid formation and the cytotoxicity of protein aggregates, which is of great significance for better understanding the pathological process and exploring the possibility of future treatment [65]. Extensive experimental and theoretical studies explored the natively stable intermediate states and complete folding pathways of the SH3 protein, and found that the unfolded state of β_1_ may be responsible for misfolding [66].

Here, we predicted the folding pathway of SH3 from Escherichia coli (eSH3, Figure 5a) and chicken c-Src-SH3 domain (cSH3, Figure 5d). Interestingly, the folding pathways and the contact order of intermediate states are almost identical for the two proteins. First, the I1 in Figure 5b and e show that a folded nucleus consisting of β_2_, β_3_, and β_4_ forms. Then, both loops marked in I2 of Figure 5b and e is called RT-Src loop, which is gradually formed. Finally, the folding nucleus is used as the support point to drive the formation of β_1_ and β_5_ at the N-terminal and C-terminal of the protein to complete the folding. The predicted results are completely consistent with the biological experimental results [67], which verifies the effectiveness of Pathfinder. Although there are differences in the intermediate state structures of the two SH3 proteins, the order of key local structures is highly consistent. Furthermore, we respectively predict the folding pathways of three homologous proteins of Escherichia coli SH3 protein mutant (PDB ID: 1srl), Caenorhabditis elegans SH3 (PDB ID: 1b07) and human Fyn SH3 (PDB ID: 5zau) in Supplement Figure 2 and 3. In these SH3, the formation of folded nucleus composed of β_2_, β_3_, and β4 can be observed, and the RT-Src loop usually only forms a rough outline. Therefore, it may be the instability of the RT-Src loop leads to insufficient constraints on β_5_ during SH3 folding, leading to misfolding. The result also show that these homologous proteins may share the same folding pathway.

**Figure 5.**
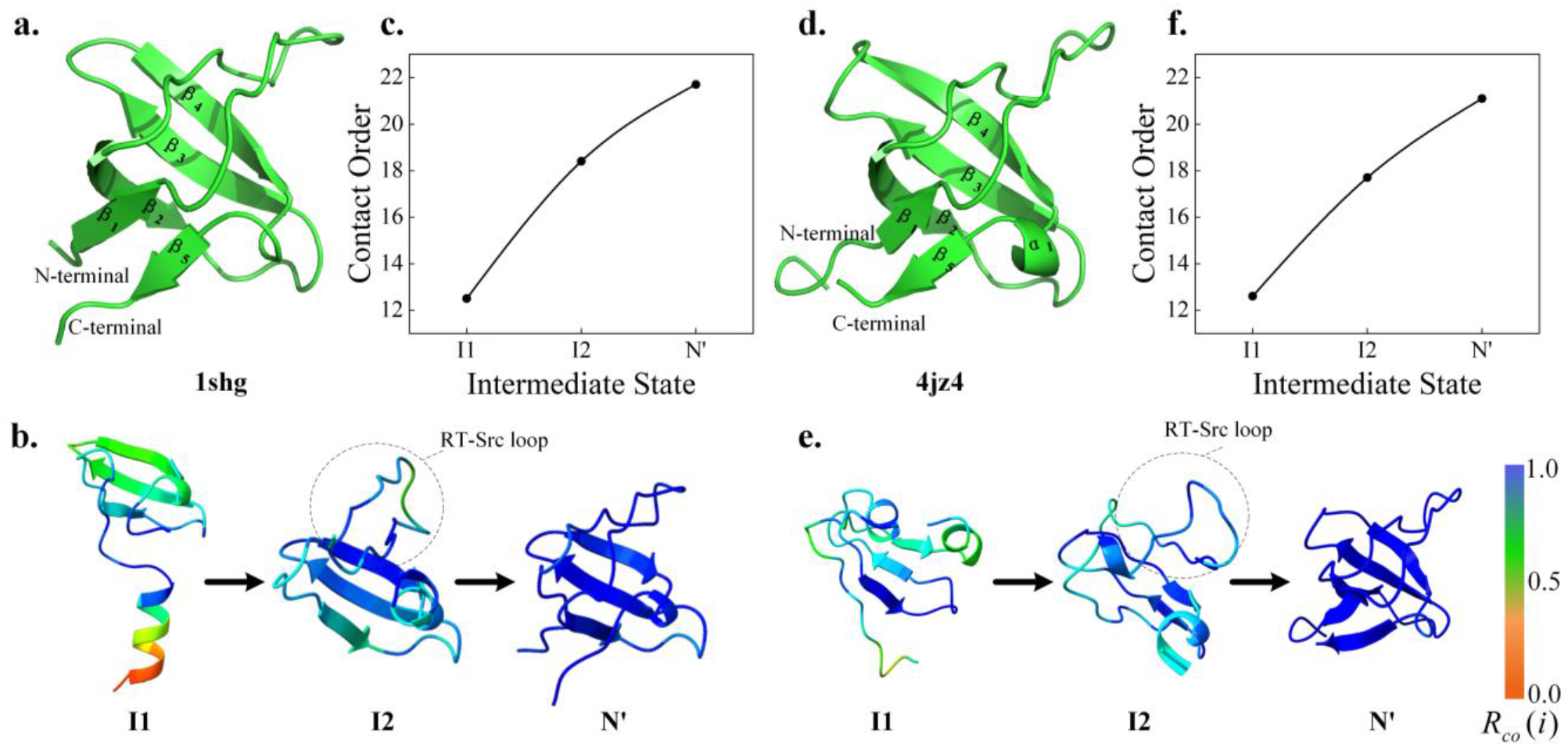
Folding pathways of eSH3 (PDB ID:1shg) and cSH3 (PDB ID:4jz4). **(a)** and **(d)** respectively their native structures. **(b)** and **(e)** are folding pathway including the intermediate states. **(c)** and **(f)** are contact order of intermediate states.

#### 3.2.3 LysM domain

The lysin domain (LysM) is a ubiquitous and versatile peptidoglycan-binding module found in bacterial proteins [68]. Because of the simple structure and important biological significance of this protein, a large number of studies have analyzed the folding transition state and folding pathway of this protein [69-71]. The protein consists of 48 residues with a secondary structure arrangement of βααβ and a highly robust folding pathway.

The folding pathway predicted by Pathfinder is shown in Figure 6c, which includes the intermediate state of only α-helix formation and the process of β-strand formation. The folding pathway of LysM is relatively clear, and the folding degree of residues can be clearly analyzed through the contact sequence of residues. The two α-helices in the middle of I1 form a folding nucleus. The inward extrusion of the two stable α-helices then drives the two ends to fold (intermediate state 2). Finally, two β-strands formation of LysM completes the folding. The result is highly consistent with biological experiments [72].

**Figure 6.**
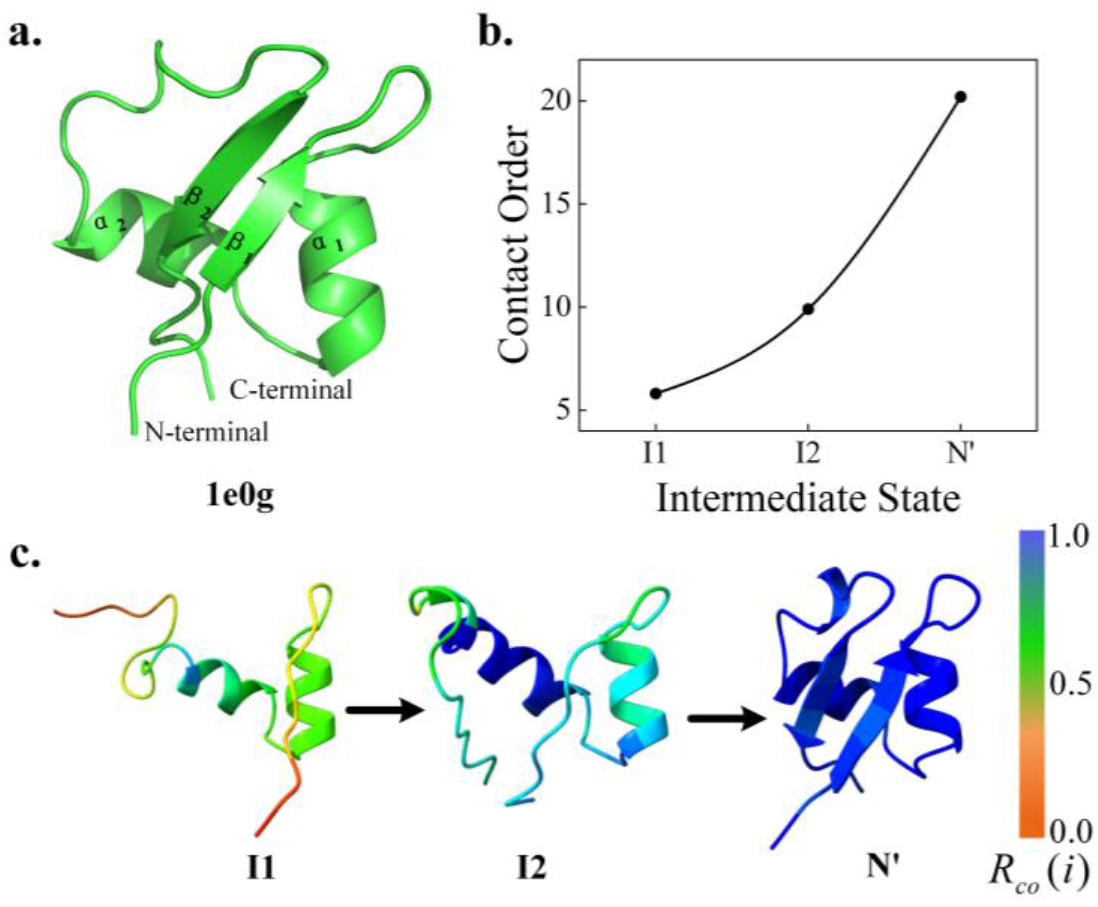
Folding pathway of LysM (PDB ID: 1e0g). **(a)** is the native structure of the LysM protein. **(b)** is the contact order of the intermediate states. **(c)** is the folding pathway including the intermediate states and their residue contact order.

### 3.3 Predicted folding pathway on proteins with partial experimental data

Because of the difficulty of obtaining the intermediate states, usually the biological experiments analyze the key residues in the folding process to study the folding mechanism. Pathfinder predicts protein heterogeneous conformation and folding pathways by sequence and native state. This may provide a new idea for the study of folding mechanism. Our predicted folding pathways for the remaining 25 proteins are presented in Supplementary Figure 2, 3 and 4. We found that for proteins with both α-helices and β-strands, the intermediate state I1 generally contains helical structures. Also, it is generally believed that the α-helical structure is more stable due to having more hydrogen bonds than the β-strand. Therefore, we think that α-helices may generally be easier to form early in folding.

Further, we classified and analyzed the results according to the contact order. As shown in Supplementary Figure 2, the increasing trend of contact order in the intermediate state of 15 proteins is the same as the case. As for Supplementary Figure 4, we found that the contact order in the intermediate state is almost higher than the native one. We further analyzed the energy force fields of 1e0m protein and 1opa protein as shown in Supplementary Figure 7. We performed a basic protein conformational sampling procedure for 1e0m protein and 1opa protein to analyze their energy force fields. The results show that the energy force field in the 1e0m protein sampling process is not accurate, and there is a situation where the energy cannot be reduced. This also shows that the appropriateness of the energy force field affects the accuracy of protein folding pathway prediction. The result of 1opa shows that the energy can be reduced normally and can be sampled to a near-natural state. However, there are cavities inside the protein’s native structure, which leads to the possibility that the residue contact order of the intermediate states is close to the native state. This also explains that in Supplementary Figure 3, there is a situation where the intermediate state is first discrete and then aggregated. Therefore, the contact order may also be deficient. In general, most of the proteins in the test set can explore the heterogeneous conformational structure information to help the understand of folding mechanism.

### 3.4 Predicted folding pathway on proteins without experimental validation

Response regulator proteins utilize distinct molecular surfaces in inactive and active conformations for various regulatory intramolecular and intermolecular protein interactions [73]. Molecular dynamics simulations complement structural studies of conformational changes under receptive domain switch function [74]. However, access to the heterogeneous conformation required for MD is not easy. Pathfinder predicted the folding pathways of four related proteins (as show in Figure 7), including a two-component response regulator from Cytophaga hutchinsonii (PDB ID: 3ilh), a response regulator from Geobacillus stearothermophilus (PDB ID: 6swl), a two-component response regulator from Clostridium difficile (PDB ID: 2qzj), and a phosphotransferase in complex with a receiver domain (PDB ID: 4qpj).

**Figure 7.**
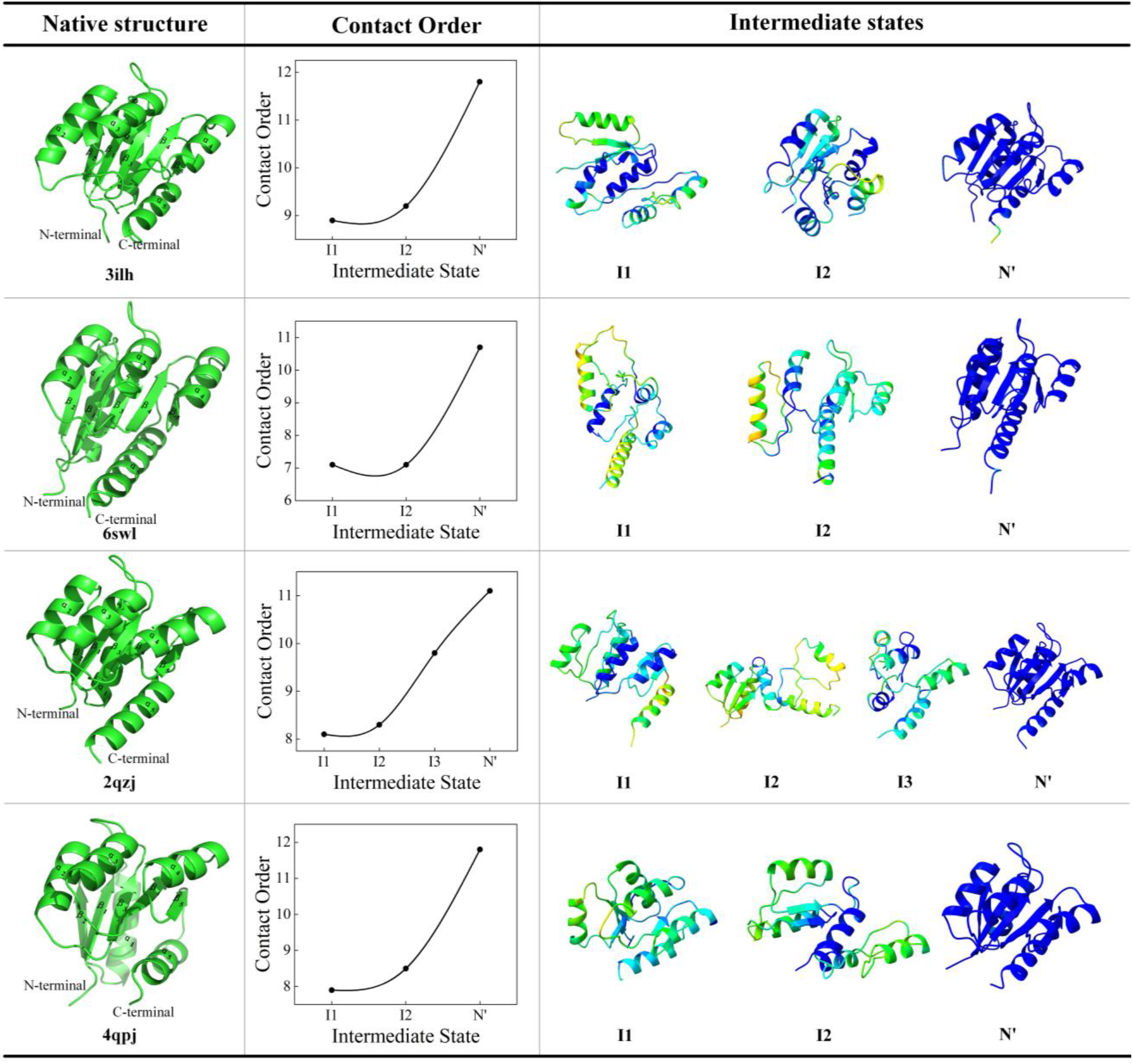
Folding pathways of four unexperimented proteins. Folding pathways include intermediate state structure information and normalized residue contact order. The color annotation of residue contact order is consistent with the intermediate state in Figure 4.

Their contact orders all increase exponentially, indicating that the intermediate conformational structure occurs more in the early stage of folding. In the early stage of folding, these proteins all form multiple α-helices, further illustrating that the helical structure may be formed earlier than the β-strand. During the subsequent folding pathway, internal β-strands are formed step by step. By comparing the intermediate states of these proteins, we found that several α-helices and β-strands often form a super-secondary structure or folded nucleus. It is the mutual extrusion and collapse of these folded nuclei that stabilize the protein fold. Pathfinder can predict the protein folding pathway by sequence and native to study the protein folding mechanism of the unresolved native conformation.

## 4. Conclusion

We apply efficient sampling algorithms to explore intermediate states and develop a protein folding pathway prediction algorithm based on conformational sampling. Pathfinder captures information between intermediate states to predict protein folding pathways. The results show that Pathfinder can extract the commonality from the folding pathways of multiple proteins, and discover the folding mechanisms of some proteins. For example, structural analogs may have different folding pathways to express different biological functions and provide insights for protein design. The proteins of the SH3 family may have the same folding pathway, and the instability of the loop region leads to insufficient force on the local structure, resulting in misfolding. During folding, α-helices may form earlier than β-strands because of the influence of hydrogen bonds. However, the experimental results show that the inaccuracy of the energy force field leads to the wrong prediction of the folding pathway. Since Pathfinder is based on the simulation of protein folding guided by the Rosetta force field. Energy force fields trained with deep learning may be applicable to folding pathway prediction for more proteins.

Compared with traditional biological experiments, Pathfinder obtains approximate protein intermediate state structures and, at the same time, predicts the order in which these intermediate states appear, enriching protein folding data. For molecular dynamics simulations, if the protein sequence is too long, the vast calculation parameters will inevitably limit the simulation of the folding process. The method can be combined with molecular dynamics simulations to provide new insights into methods for computationally simulating protein folding. At the same time, Pathfinder can analyze family proteins or protein collections under different classifications, and explore protein folding mechanisms from a broader perspective.

## Key points

- We modify the energy function to make conformational sampling explore the transition tendency of intermediate states, which is a valuable attempt to explore intermediate states through conformational sampling algorithms.
- We propose a protein folding pathway prediction algorithm (Pathfinder). Pathfinder further obtained the structural information of the intermediate state by large-scale sampling, combined with the above algorithm to explore the transition probability of the intermediate state, and predicted the protein folding pathway.
- The results of the dataset demonstrate the effectiveness of Pathfinder in predicting protein folding pathways. The results also reveal folding mechanisms, such as that structural analogs may fold differently to express different biological functions, homologous proteins may share the same folding pathway, and α-helices may form earlier in folding than β-strands.

## Data availability

All data needed to evaluate the conclusions are present in the paper and the Supplementary Materials. The folding data in this paper and web server of Pathfinder is freely available at http://zhanglab-bioinf.com/Pathfinder/.

## Supplementary Information

### Supplementary Figure

**Figure 1.**
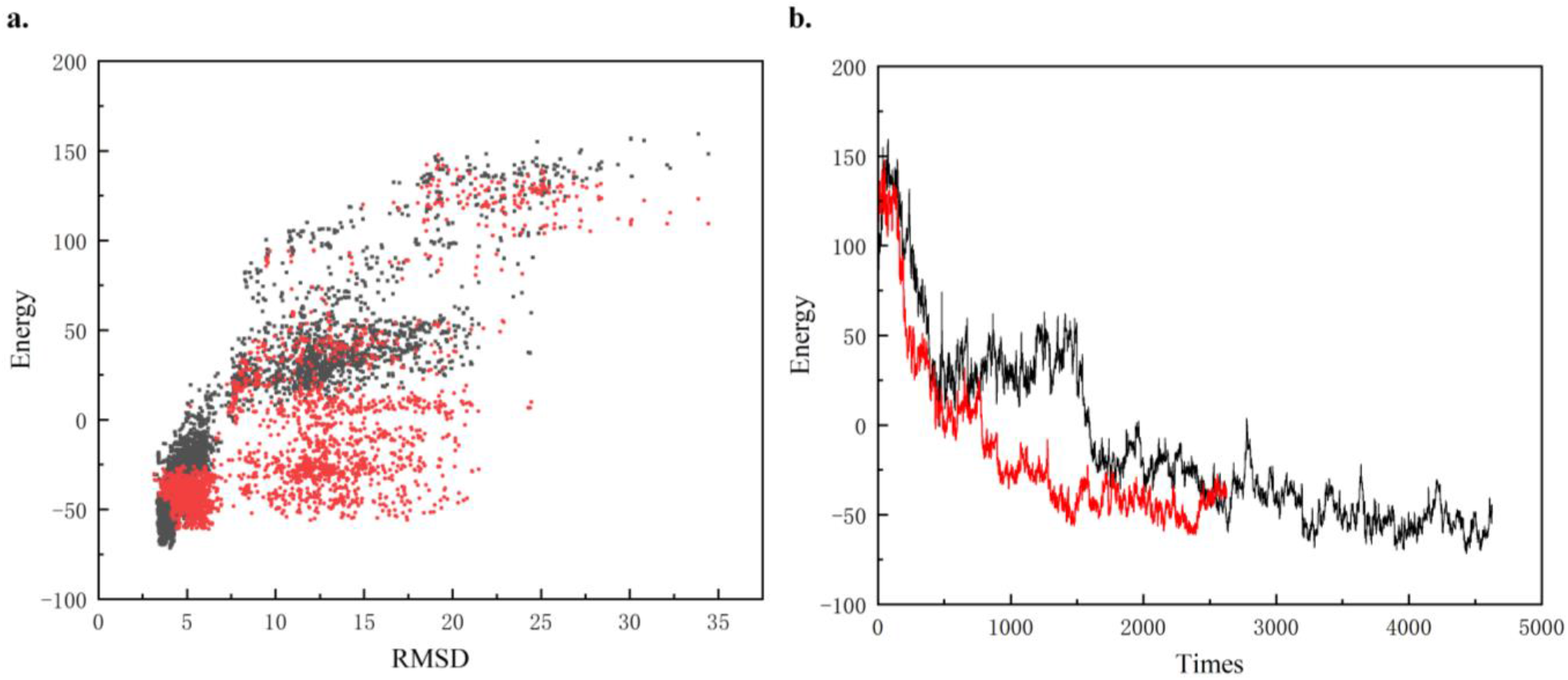
**(a)** is the energy and RMSD scatterplot of the conformations of the sampling process. Red dots are the conformations of the sampling process under the modified energy force field. Black dots are the benchmark conformations of Rosetta’s conformational sampling process. **(b)** is the trajectory of energy variation with the number of samples. Red is the sampling trace of the modified energy force field, and black is the sampling trace of the baseline Rosetta.

We apply the modified energy function to Rosetta’s ClassicAbinitio protocol and compare the accuracy of the structure prediction with the baseline’s version. Since the conformational sampling algorithm stops the conformational sampling process when the energy converges, we count the sampling times from conformational sampling to convergence to compare the sampling efficiency. The results of 331 tested proteins showed that the modified energy function reduced the number of samples by 12.5% without losing the accuracy of the predicted protein. A summary table is shown in Supplementary Table 3.

We further analyzed the energy changes of the conformational sampling process, as shown in Supplementary Figure 1 for the comparison of the conformational sampling process of 2qzj_A protein. The prediction accuracy of the 2qzj_A protein improved from 0.5215 of the baselines to 0.6799, while sampling faster to convergence as shown in Supplementary Fig 1b. The results show that the modified energy function can improve the sampling efficiency and make it easier to explore the energy basin to obtain the state transition tendency.

**Figure 2.**
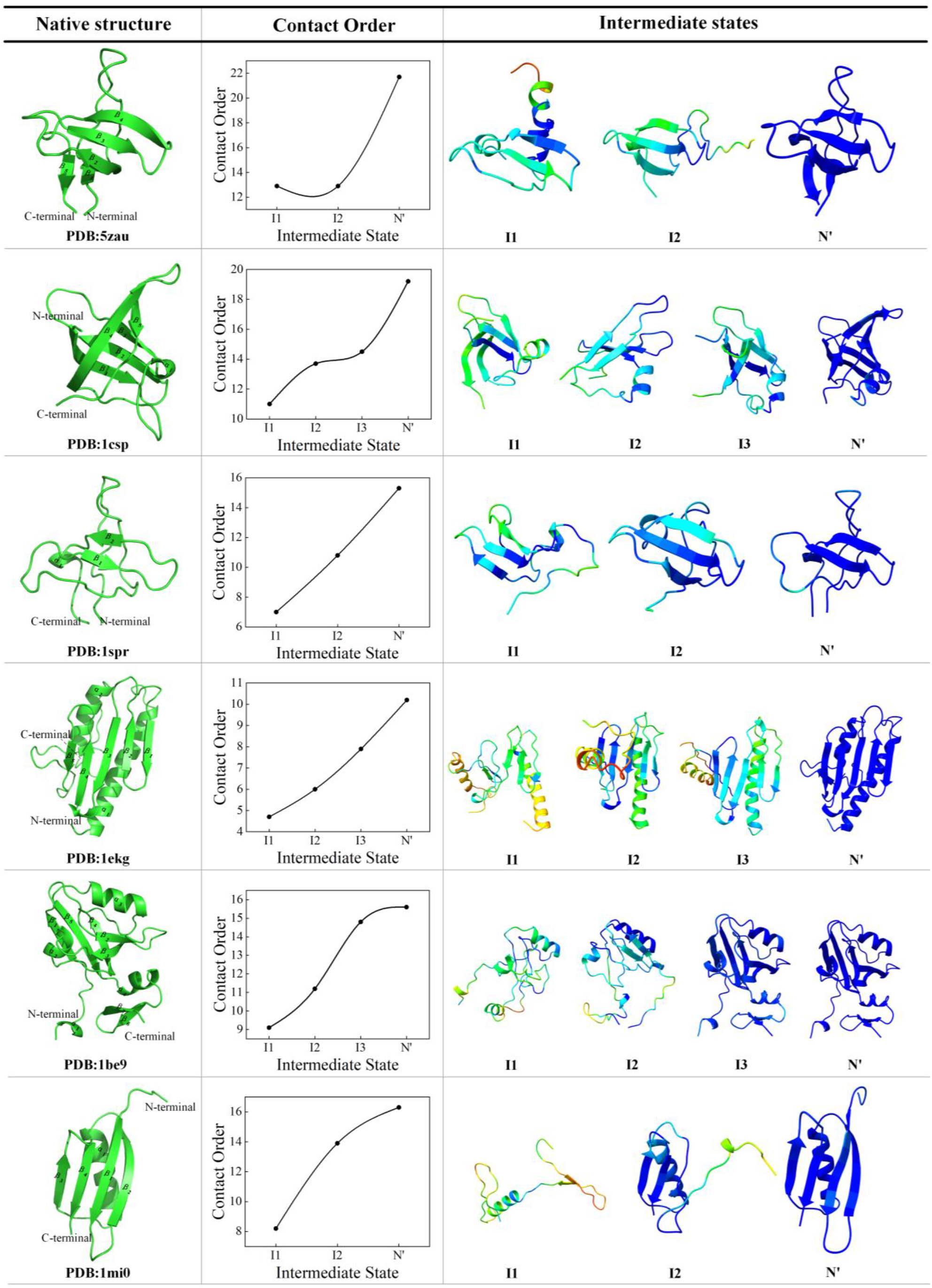
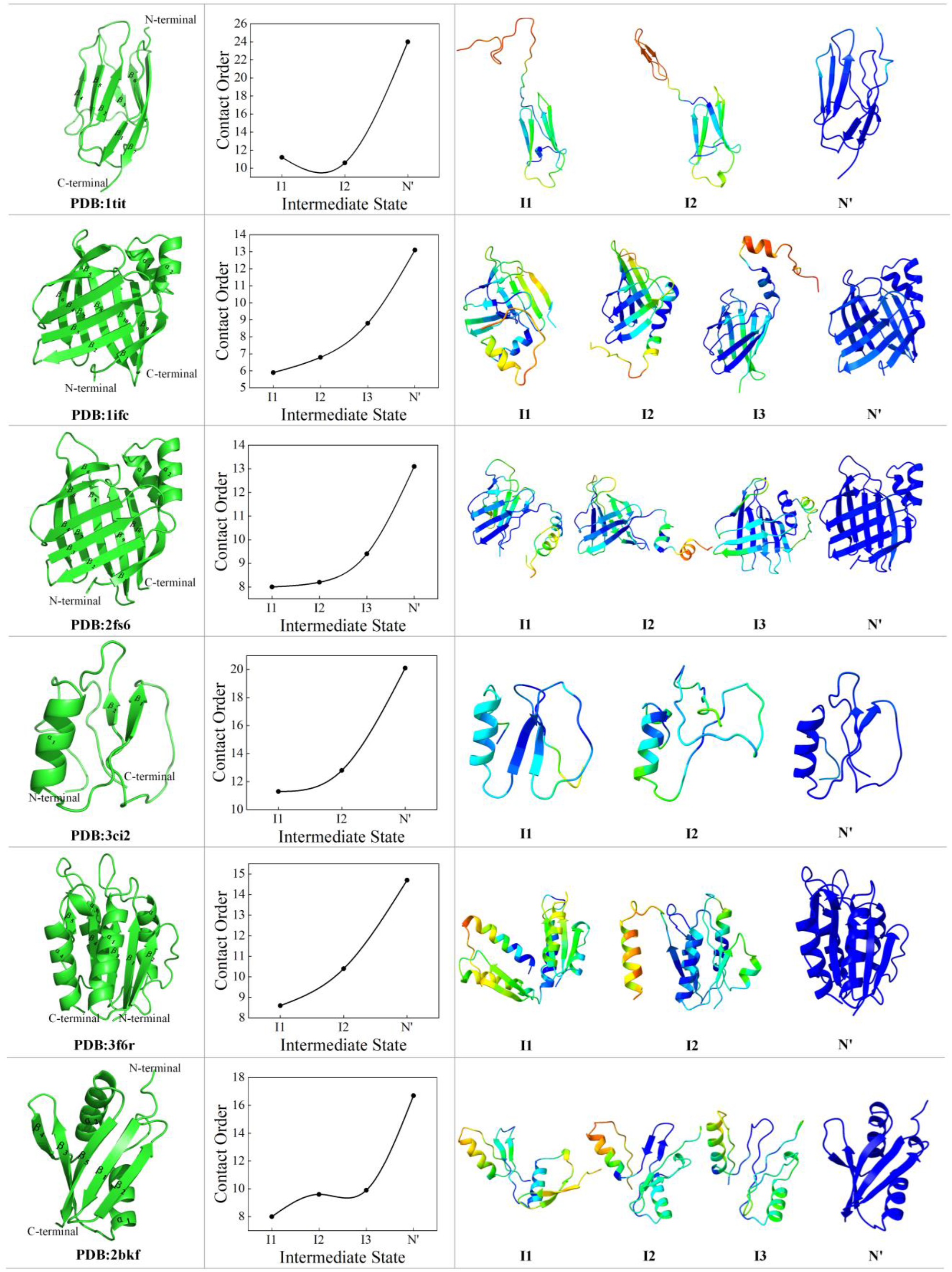
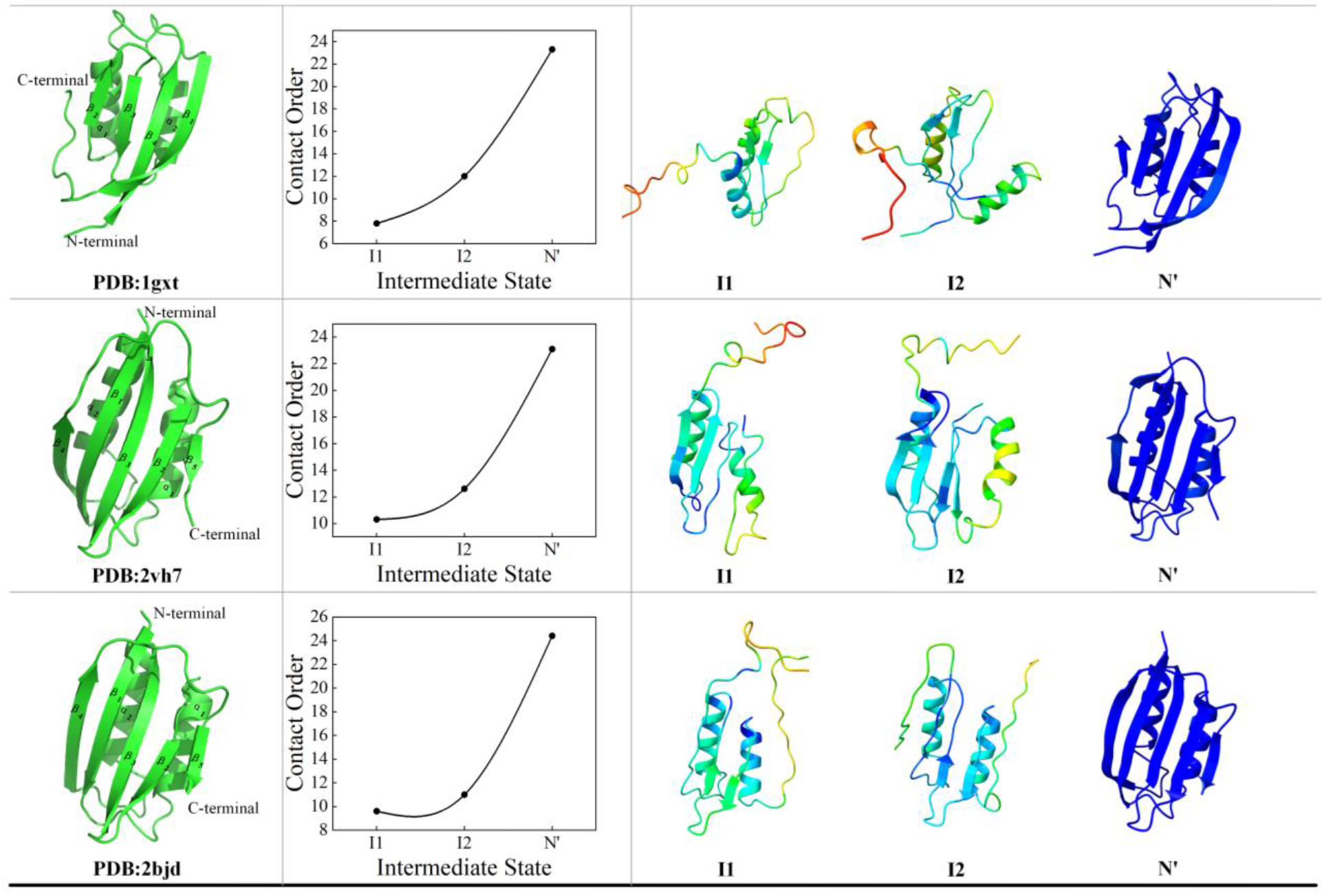
Folding pathways of 15 proteins that the contact order shows an increasing trend.

**Figure 3.**
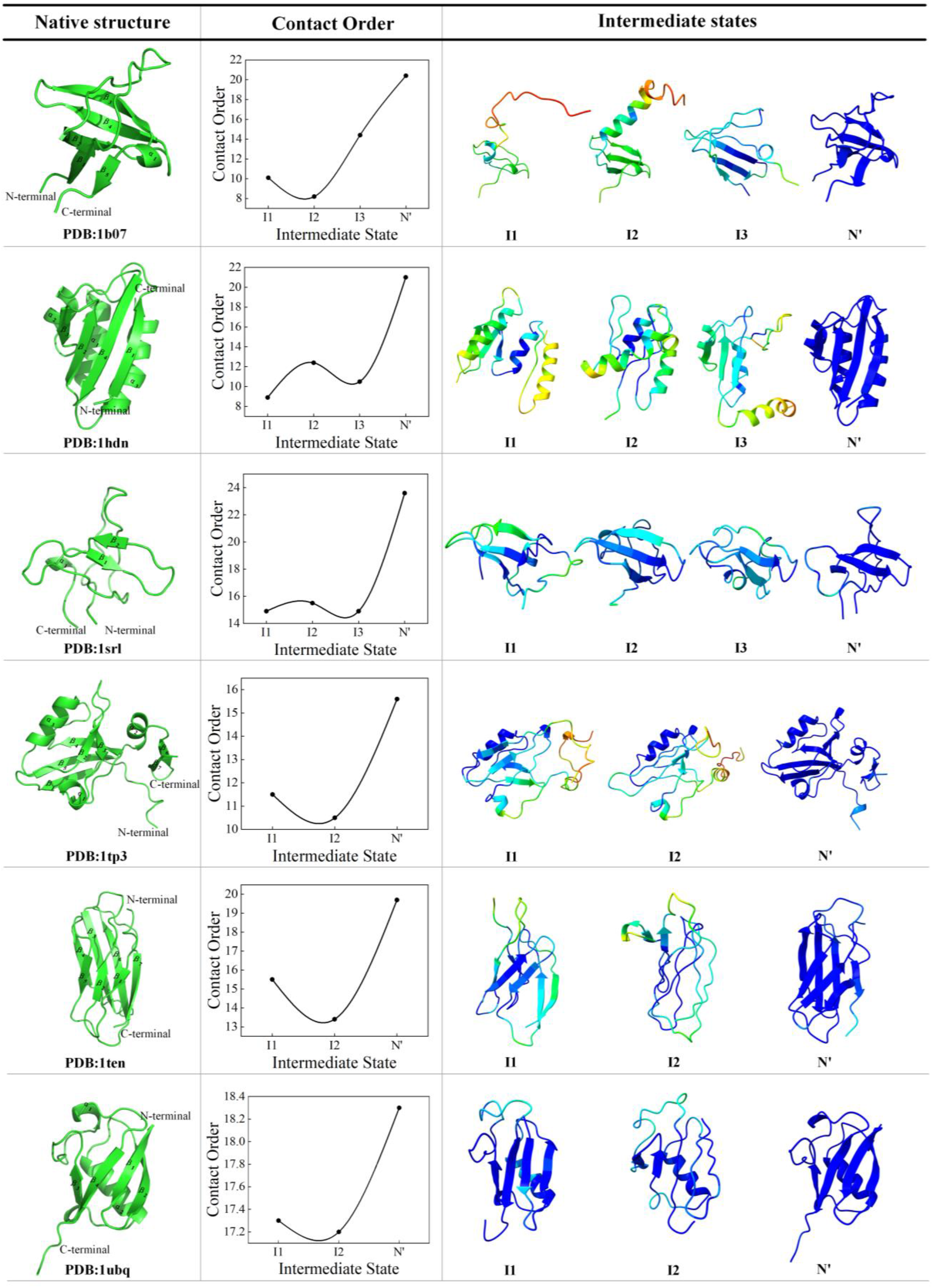
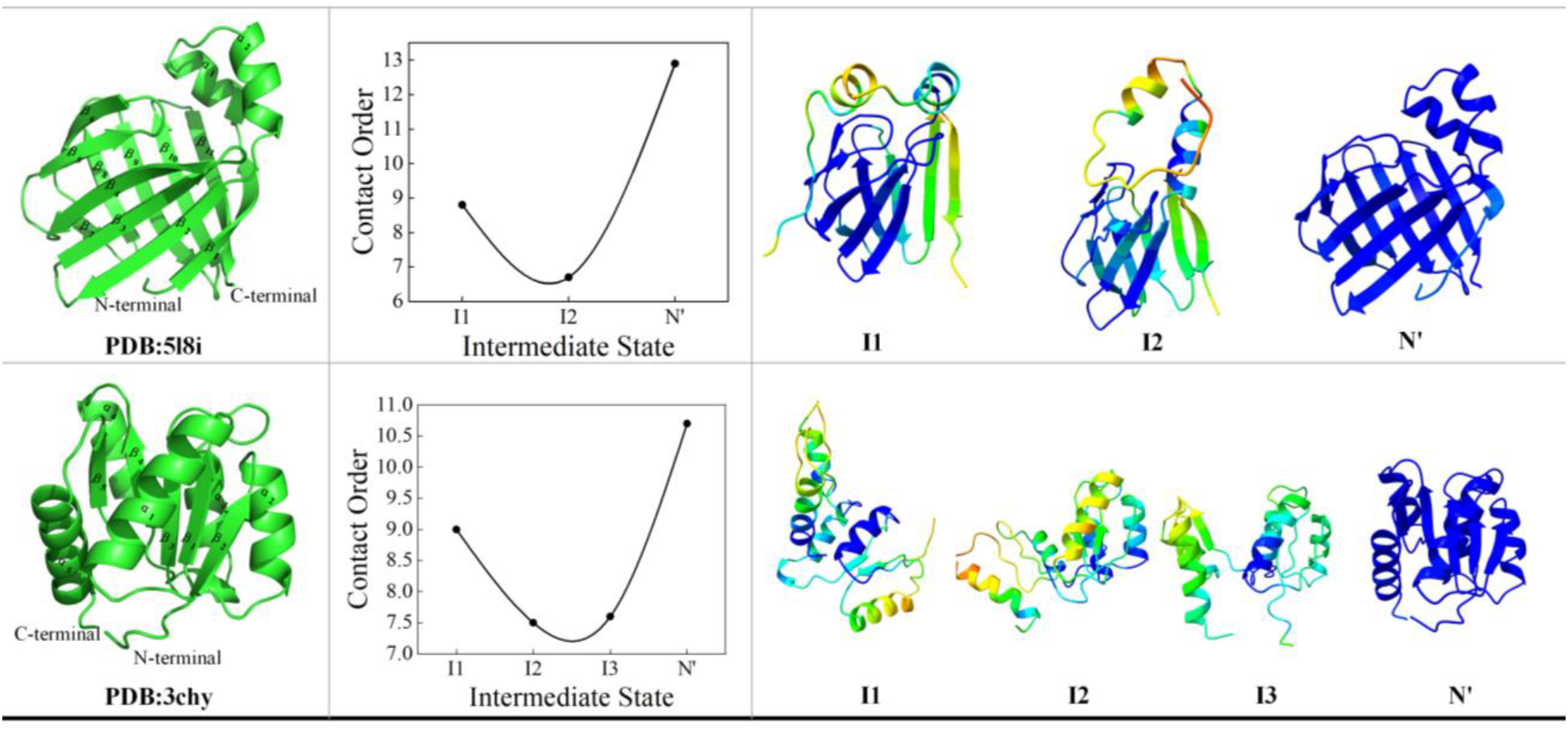
Folding pathways of 8 proteins that include intermediate states.

**Figure 4.**
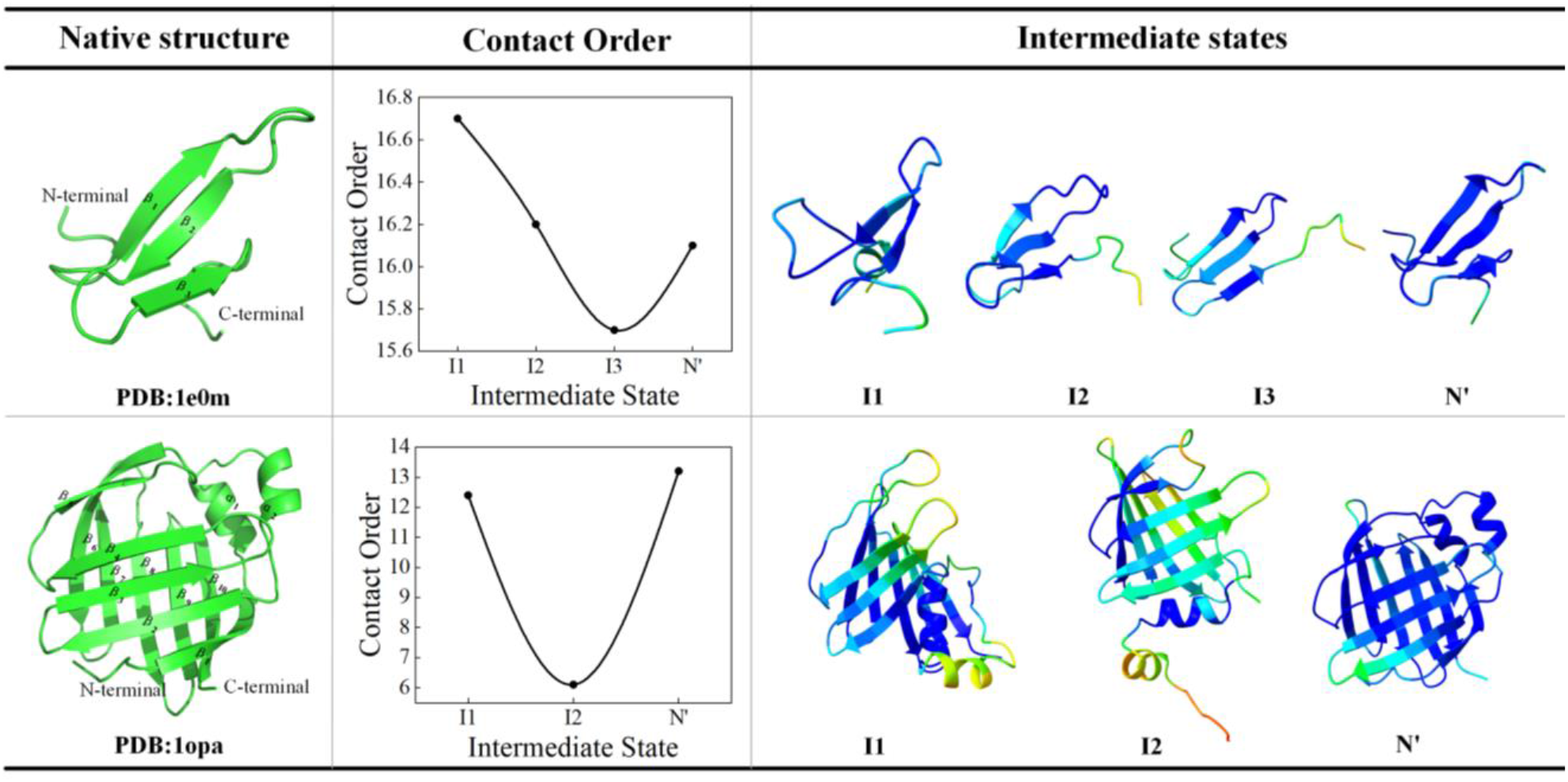
The folding pathway of 2 proteins with bias contact order.

**Figure 5.**
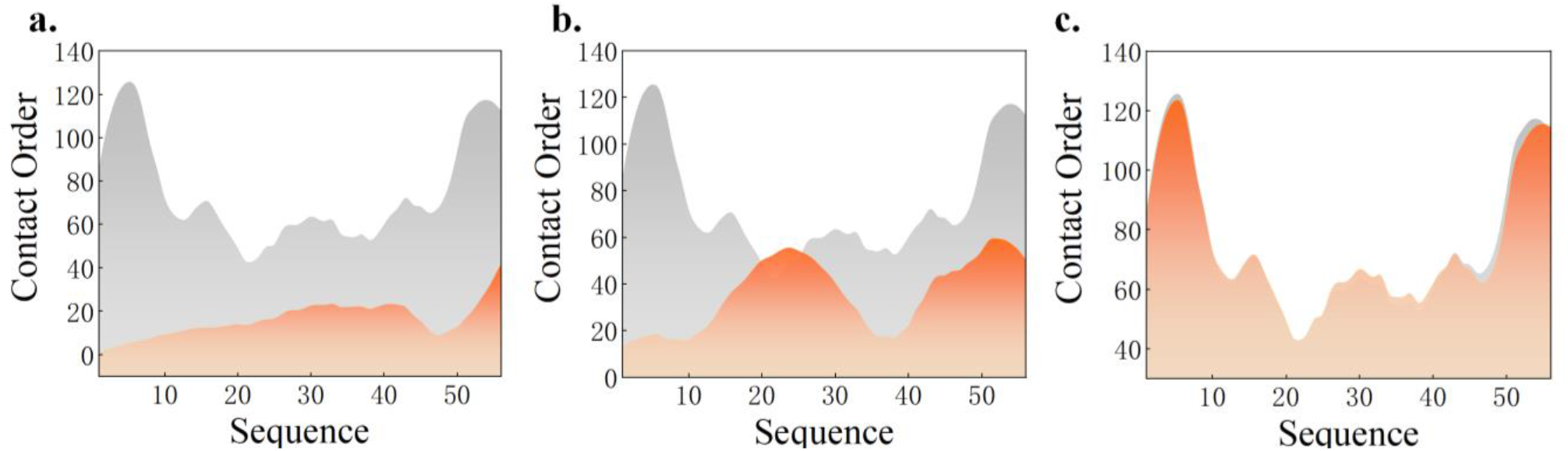
The spline connection graph of the normalized residue contact order. The gray is native state and the orange is intermediate state, both of which come from the B1 domain of protein G.

**Figure 6.**
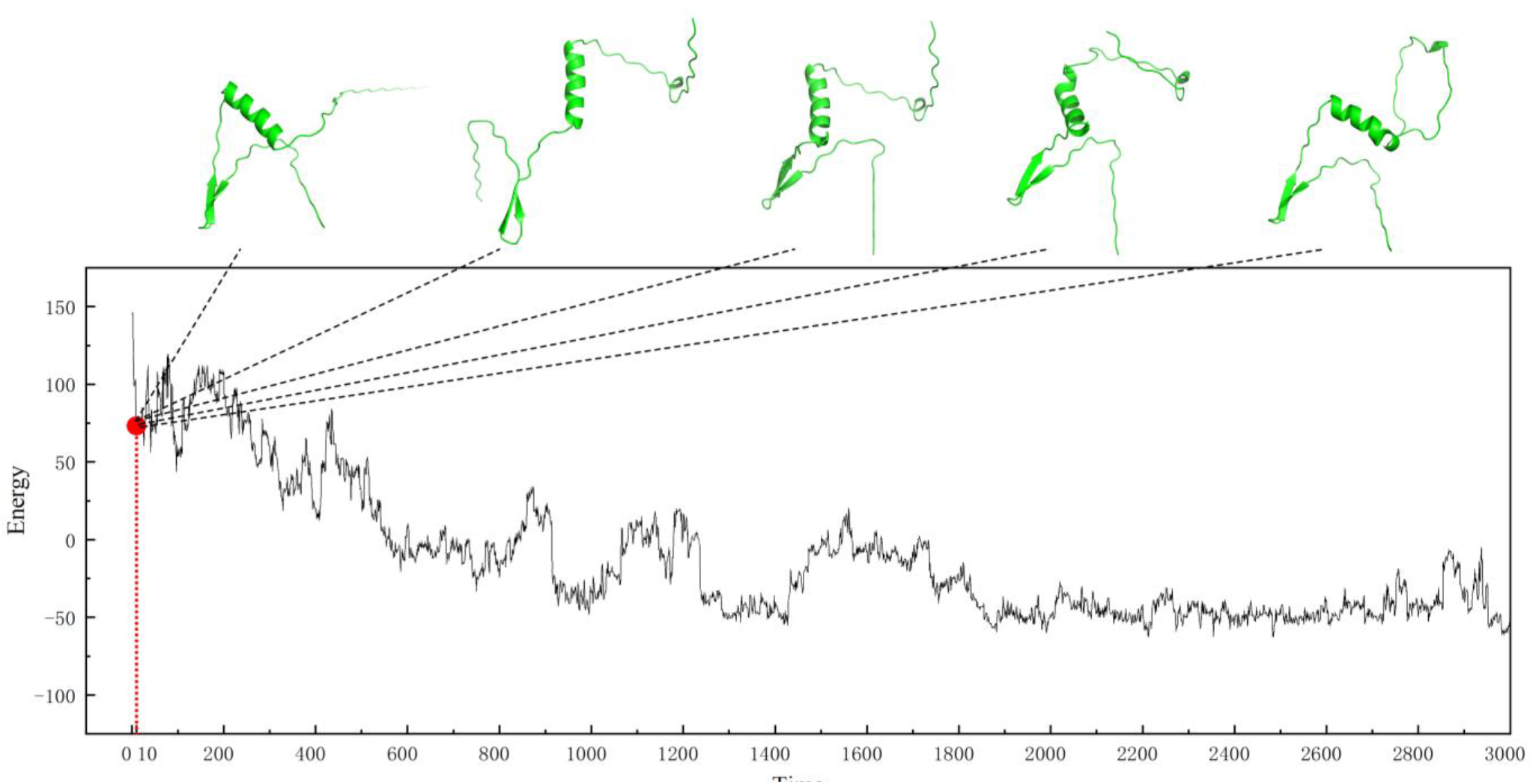
LB1 sampling process diagram. A total of 10125 accepted process points were generated during one conformational sampling process of LB1. We gave the first 3000 sampling data and analyzed the first 30 conformations. And present part of the conformation.

**Figure 7.**
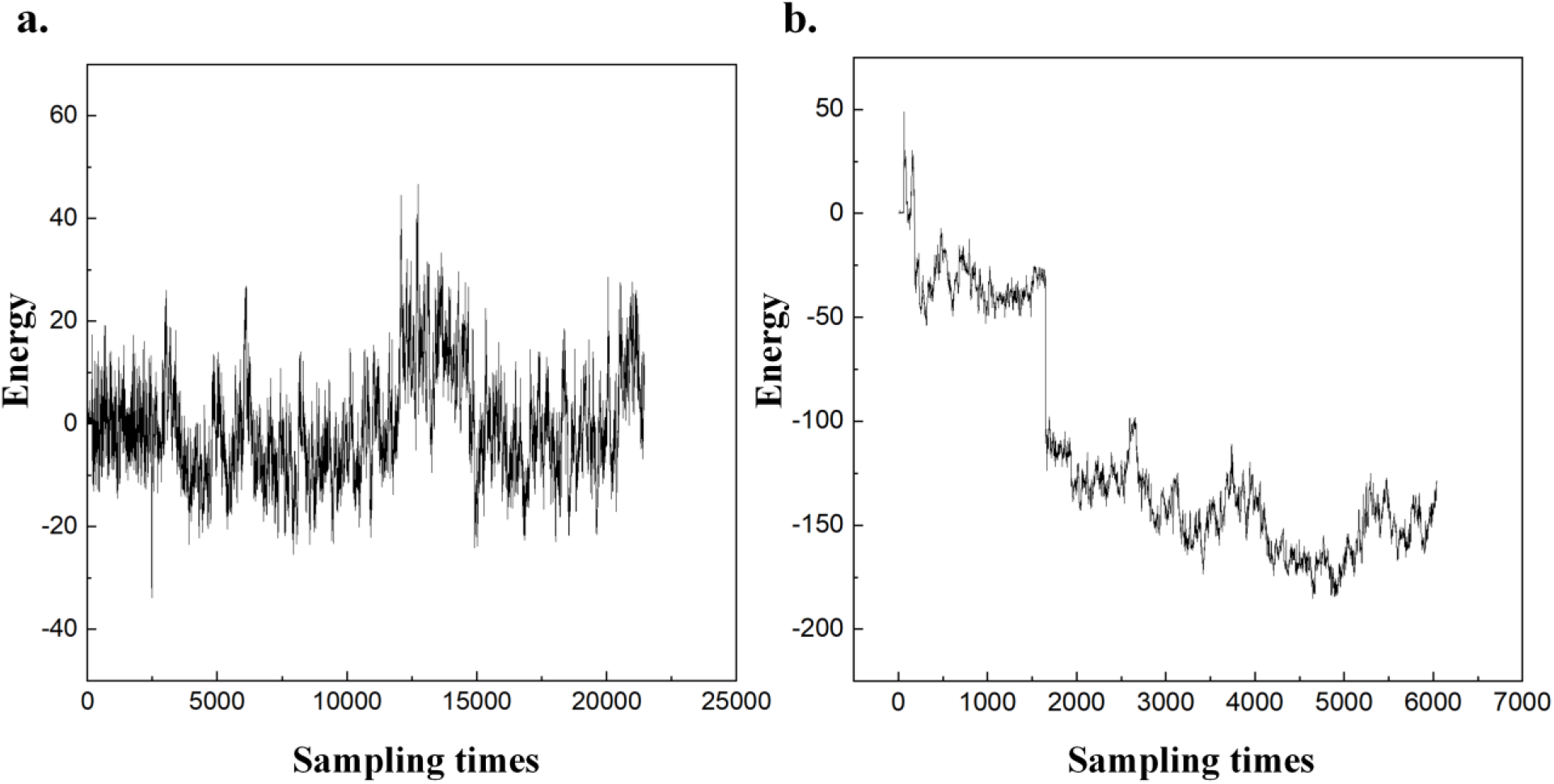
Line graph of sampling times and energy. **(a)** is the sampling trajectory of 1e0m protein, and **(b)** is the sampling trajectory of 1opa. Among them, the protein energy of 1e0m has almost no drop, and 1opa is more in line with the normal conformational sampling process.

### Supplementary Tables

**Table S1.**
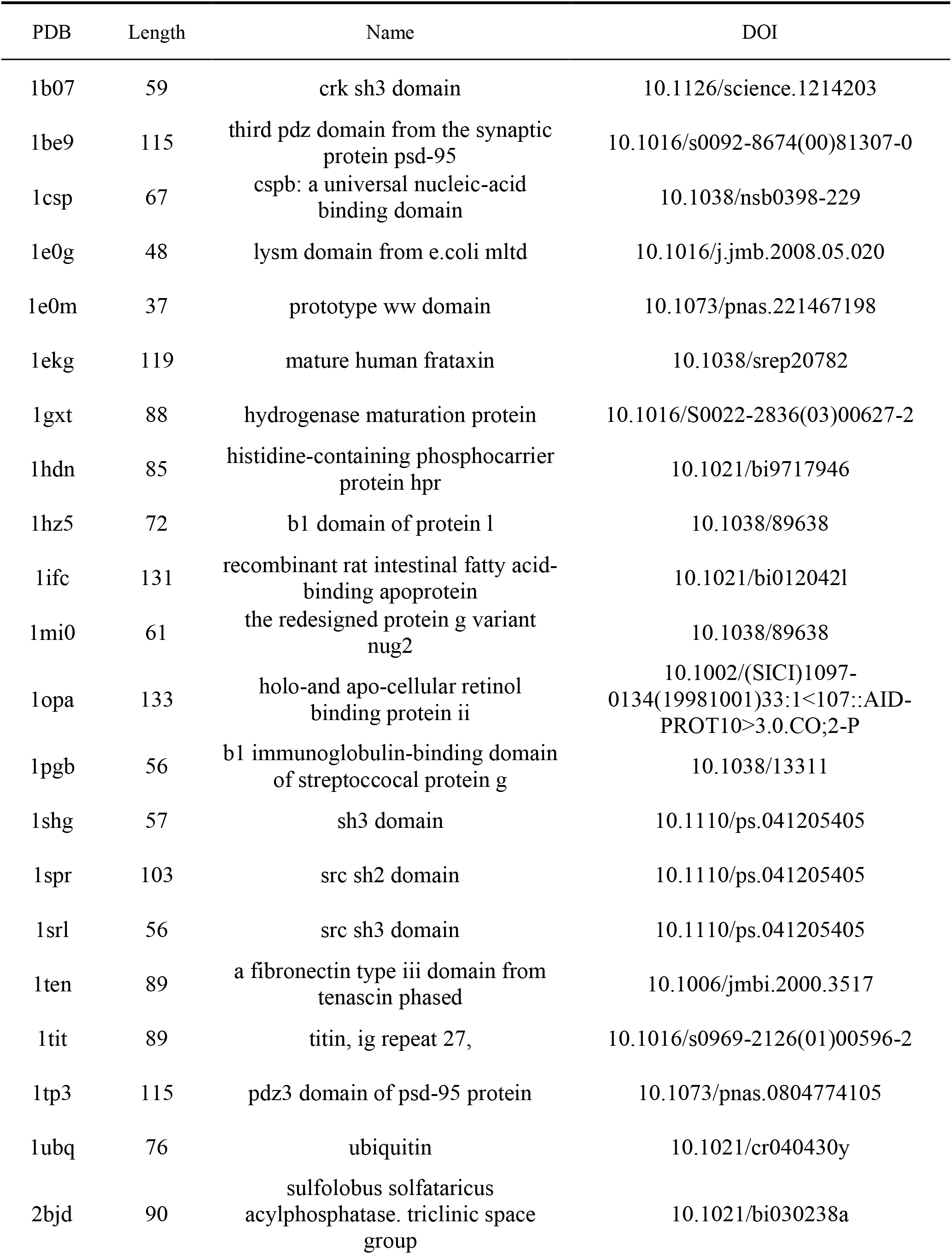
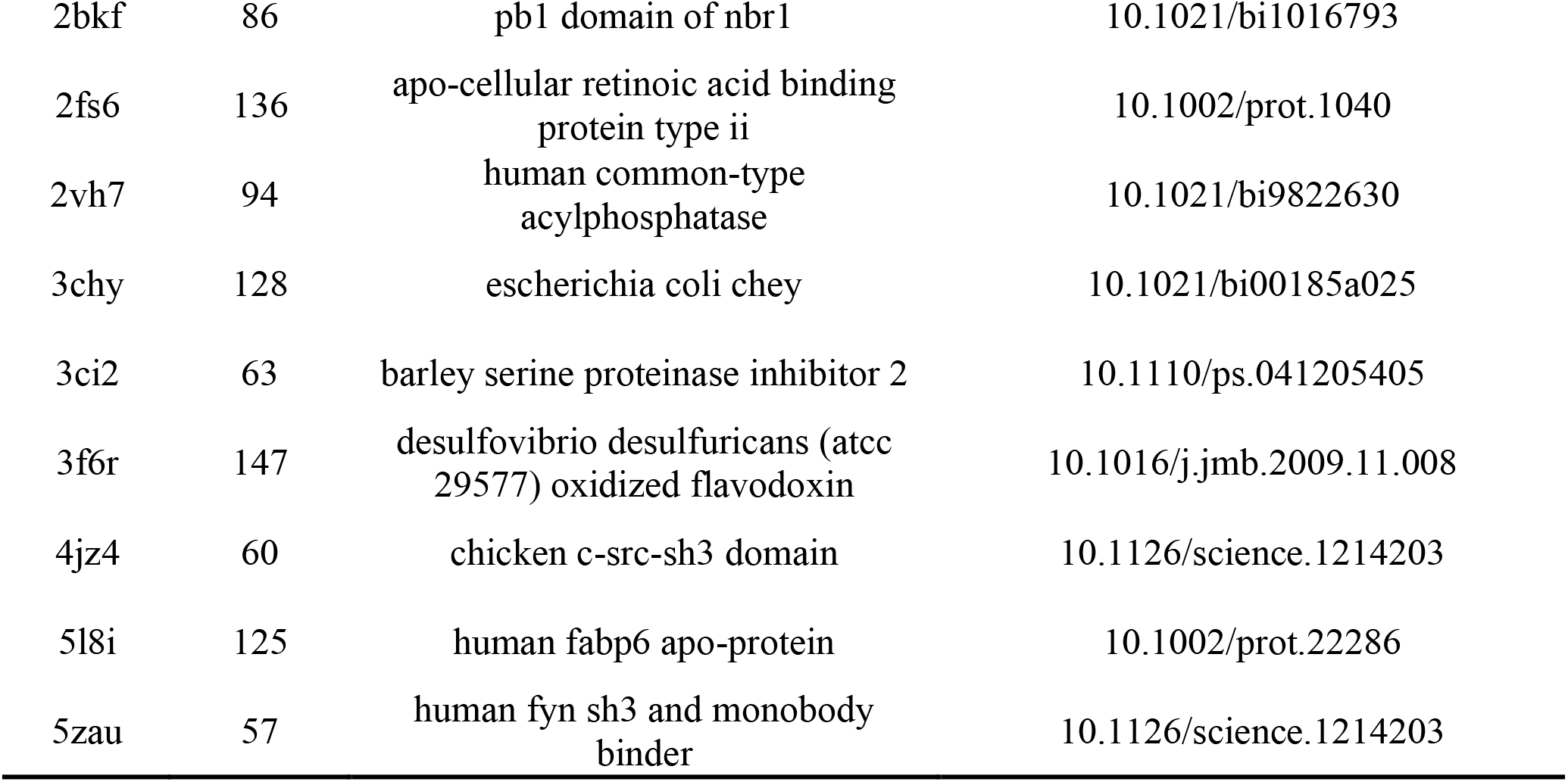
The dataset of Pathfinder.

**Table S2.** The parameter of Pathfinder.

| Parameter | Default Value |
| --- | --- |
| $G$ | 10 |
| $S$ | 10 |
| $\tau$ | 0.55 |
| $\mu$ | 1000 |
| $\eta$ | 0.5 |
| $\xi$ | 0.7 |

**Table 3-1.** Performance improvement table of modified energy function (MEF)

| Methods | TM-score | Sample_time | Running_time(min) |
| --- | --- | --- | --- |
| MEF | **0.4258** | **23813** | **185** |
| Rosetta | 0.4254 | 27227 | 193 |

## References

[1]. Jumper J, Evans R, Pritzel A et al. Highly accurate protein structure prediction with AlphaFold, Nature 2021;596:583–589.

[2]. Baek M, DiMaio F, Anishchenko I et al. Accurate prediction of protein structures and interactions using a three-track neural network, Science 2021;373:871-+.

[3]. AlQuraishi M. Machine learning in protein structure prediction, Current Opinion in Chemical Biology 2021;65:1–8.

[4]. Varadi M, Anyango S, Deshpande M et al. AlphaFold Protein Structure Database: massively expanding the structural coverage of protein-sequence space with high-accuracy models, Nucleic Acids Research 2022;50:D439–D444.

[5]. Callaway E. ’the Entire Protein Universe’: Ai Predicts Shape of Nearly Every Known Protein, Nature 2022;608:15–16.

[6]. Service RF. STRUCTURAL BIOLOGY ‘The game has changed.’ AI triumphs at protein folding, Science 2020;370:1144–1145.

[7]. Moore PB, Hendrickson WA, Henderson R et al. The protein-folding problem: Not yet solved, Science 2022;375:507–507.

[8]. Chen SJ, Hassan M, Jernigan RL et al. Protein folds vs. protein folding: Differing questions, different challenges, Proceedings of the National Academy of Sciences 2023;120:e2214423119.

[9]. Jones DT, Thornton JM. The impact of AlphaFold2 one year on, Nature Methods 2022;19:15–20.

[10]. Fowler NJ, Williamson MP. The accuracy of protein structures in solution determined by AlphaFold and NMR, Structure 2022;30:925–933. e922.

[11]. Outeiral C, Nissley DA, Deane CM et al. Current structure predictors are not learning the physics of protein folding, Bioinformatics 2022;38:1881–1887.

[12]. Stiller JB, Otten R, Haussinger D et al. Structure determination of high-energy states in a dynamic protein ensemble, Nature 2022;603:528–535.

[13]. Kopito RR. Aggresomes, inclusion bodies and protein aggregation, Trends in Cell Biology 2000;10:524–530.

[14]. Song J, Takemoto K, Shen H et al. Prediction of Protein Folding Rates from Structural Topology and Complex Network Properties, IPSJ Transactions on Bioinformatics 2010;3:40–53.

[15]. Valastyan JS, Lindquist S. Mechanisms of protein-folding diseases at a glance, Disease Models & Mechanisms 2014;7:9–14.

[16]. Selkoe DJ, Hardy J. The amyloid hypothesis of Alzheimer’s disease at 25 years, EMBO Molecular Medicine 2016;8:595–608.

[17]. Kalia LV, Kalia SK, Lang AE. Disease-Modifying Strategies for Parkinson’s Disease, Movement Disorders 2015;30:1442–1450.

[18]. Hartl FU. Protein Misfolding Diseases, Annual Review of Biochemistry 2017;86:21–26.

[19]. Baker D. What has de novo protein design taught us about protein folding and biophysics?, Protein Science 2019;28:678–683.

[20]. Huang PS, Boyken SE, Baker D. The coming of age of de novo protein design, Nature 2016;537:320–327.

[21]. Ni D, Chai Z, Wang Y et al. Along the allostery stream: Recent advances in computational methods for allosteric drug discovery, Wiley Interdisciplinary Reviews: Computational Molecular Science 2022;12:e1585.

[22]. Direito I, Fardilha M, Helguero LA. Contribution of the unfolded protein response to breast and prostate tissue homeostasis and its significance to cancer endocrine response, Carcinogenesis 2019;40:203–215.

[23]. Chiti F, Dobson CM. Protein Misfolding, Amyloid Formation, and Human Disease: A Summary of Progress Over the Last Decade, Annual Review of Biochemistry 2017;86:27–68.

[24]. Neudecker P, Robustelli P, Cavalli A et al. Structure of an Intermediate State in Protein Folding and Aggregation, Science 2012;336:362–366.

[25]. Guinn EJ, Jagannathan B, Marqusee S. Single-molecule chemo-mechanical unfolding reveals multiple transition state barriers in a small single-domain protein, Nature Communications 2015;6:6861.

[26]. Bhatia S, Udgaonkar JB. Heterogeneity in Protein Folding and Unfolding Reactions, Chemical Reviews 2022;122:8911–8935.

[27]. Choi HK, Min D, Kang H et al. Watching helical membrane proteins fold reveals a common N-to-C-terminal folding pathway, Science 2019;366:1150–1156.

[28]. Korzhnev DM, Kay LE. Probing invisible, low-populated states of protein molecules by relaxation dispersion NMR spectroscopy: An application to protein folding, Accounts of Chemical Research 2008;41:442–451.

[29]. Baldwin RL. The search for folding intermediates and the mechanism of protein folding, Annual Review of Biophysics 2008;37:1–21.

[30]. Maxwell KL, Wildes D, Zarrine-Afsar A et al. Protein folding: Defining a "standard" set of experimental conditions and a preliminary kinetic data set of two-state proteins, Protein Science 2005;14:602–616.

[31]. Feng H, Zhou Z, Bai Y. A protein folding pathway with multiple folding intermediates at atomic resolution, Proceedings of the National Academy of Sciences 2005;102:5026–5031.

[32]. Hong H, Guo Z, Sun H et al. Two energy barriers and a transient intermediate state determine the unfolding and folding dynamics of cold shock protein, Communications Chemistry 2021;4:156.

[33]. Masson GR, Burke JE, Ahn NG et al. Recommendations for performing, interpreting and reporting hydrogen deuterium exchange mass spectrometry (HDX-MS) experiments, Nature Methods 2019;16:595–602.

[34]. Greenfield NJ. Analysis of the kinetics of folding of proteins and peptides using circular dichroism, Nature Protocols 2006;1:2891–2899.

[35]. Finkelstein AV. 50+Years of Protein Folding, Biochemistry(Moscow) 2018;83:S3–S18.

[36]. Auer S, Miller MA, Krivov SV et al. Importance of metastable states in the free energy landscapes of polypeptide chains, Physical Review Letters 2007;99:178104.

[37]. Freddolino PL, Harrison CB, Liu Y et al. Challenges in protein-folding simulations, Nature Physics 2010;6:751–758.

[38]. Lindorff-Larsen K, Piana S, Dror RO et al. How Fast-Folding Proteins Fold, Science 2011;334:517–520.

[39]. Lane TJ, Bowman GR, Beauchamp K et al. Markov State Model Reveals Folding and Functional Dynamics in Ultra-Long MD Trajectories, Journal of the American Chemical Society 2011;133:18413–18419.

[40]. Noe F, De Fabritiis G, Clementi C. Machine learning for protein folding and dynamics, Current Opinion in Structural Biology 2020;60:77–84.

[41]. Ramaswamy VK, Musson SC, Willcocks CG et al. Deep Learning Protein Conformational Space with Convolutions and Latent Interpolations, Physical Review X 2021;11:011052.

[42]. Nijhawan AK, Chan AM, Hsu DJ et al. Resolving Dynamics in the Ensemble: Finding Paths through Intermediate States and Disordered Protein Structures, Journal of Physical Chemistry B 2021;125:12401–12412.

[43]. Zhao K, Xia Y, Zhang F et al. Protein structure and folding pathway prediction based on remote homologs recognition using PAthreader, Communications Biology 2023;6:243.

[44]. Berman HM. The Protein Data Bank, Nucleic Acids Research 2000;28:235–242.

[45]. Ulmschneider JP, Ulmschneider MB, Di Nola A. Monte Carlo vs molecular dynamics for all-atom polypeptide folding simulations, Journal of Physical Chemistry B 2006;110:16733–16742.

[46]. Kmiecik S, Kolinski A. Folding pathway of the B1 domain of protein G explored by multiscale modeling, Biophysical Journal 2008;94:726-736.

[47]. Kmiecik S, Kolinski A. Characterization of protein-folding pathways by reduced-space modeling, Proceedings of the National Academy of Sciences 2007;104:12330–12335.

[48]. Zhao K, Liu J, Zhou X et al. MMpred: a distance-assisted multimodal conformation sampling for de novo protein structure prediction, Bioinformatics 2021;37:4350–4356.

[49]. Xia Y, Peng C, Zhou X et al. A sequential niche multimodal conformational sampling algorithm for protein structure prediction, Bioinformatics 2021;37:4357–4365.

[50]. Englander SW, Mayne L. The case for defined protein folding pathways, Proceedings of the National Academy of Sciences 2017;114:8253–8258.

[51]. Das R, Baker D. Macromolecular modeling with Rosetta, Annual Review of Biochemistry 2008;77:363–382.

[52]. Rohl CA, Strauss CEM, Misura KMS et al. Protein Structure Prediction Using Rosetta. Methods in Enzymology. Elsevier, 2004, 66–93.

[53]. Metropolis N, Rosenbluth AW, Rosenbluth MN et al. Equation of State Calculations by Fast Computing Machines, The Journal of Chemical Physics 1953;21:1087–1092.

[54]. Zhang Y, Skolnick J. SPICKER: A clustering approach to identify near-native protein folds, Journal of Computational Chemistry 2004;25:865–871.

[55]. Wang J, Panagiotou E. The protein folding rate and the geometry and topology of the native state, Scientific Reports 2022;12:6384.

[56]. Plaxco KW, Simons KT, Baker D. Contact order, transition state placement and the refolding rates of single domain proteins, Journal of Molecular Biology 1998;277:985–994.

[57]. Dinner AR, Karplus M. The roles of stability and contact order in determining protein folding rates, Nature Structural Biology 2001;8:21–22.

[58]. Manavalan B, Kuwajima K, Lee J. PFDB: A standardized protein folding database with temperature correction, Scientific Reports 2019;9:1–9.

[59]. Pancsa R, Varadi M, Tompa P et al. Start2Fold: a database of hydrogen/deuterium exchange data on protein folding and stability, Nucleic Acids Research 2016;44:D429–D434.

[60]. Gallagher T, Alexander P, Bryan P et al. Two Crystal Structures of the B1 Immunoglobulin-Binding Domain of Streptococcal Protein G and Comparison with NMR, Biochemistry 2002;33:4721–4729.

[61]. Park S-H, Shastry M, Roder H. Folding dynamics of the B1 domain of protein G explored by ultrarapid mixing, Nature Structural Biology 1999;6:943–947.

[62]. Chang LW, Perez A. Deciphering the Folding Mechanism of Proteins G and L and Their Mutants, Journal of the American Chemical Society 2022;144:14668–14677.

[63]. Gronenborn AM, Filpula DR, Essig NZ, et al. A Novel, Highly Stable Fold of the Immunoglobulin Binding Domain of Streptococcal Protein-G, Science 1991;253:657–661.

[64]. Nauli S, Kuhlman B, Baker D. Computer-based redesign of a protein folding pathway, Nature Structural Biology 2001;8:602–605.

[65]. Ventura S, Zurdo J, Narayanan S et al. Short amino acid stretches can mediate amyloid formation in globular proteins: The Src homology 3 (SH3) case, Proceedings of the National Academy of Sciences 2004;101:7258–7263.

[66]. Bacarizo J, Martinez-Rodriguez S, Martin-Garcia JM et al. Electrostatic Effects in the Folding of the SH3 Domain of the c-Src Tyrosine Kinase: pH-Dependence in 3D-Domain Swapping and Amyloid Formation, PloS One 2014;9:e113224.

[67]. Petzold K, Öhman A, Backman L. Folding of the αΙΙ-spectrin SH3 domain under physiological salt conditions, Archives of Biochemistry and Biophysics 2008;474:39–47.

[68]. Bateman A, Bycroft M. The structure of a LysM domain from E-coli membrane-bound lytic murein transglycosylase D (MltD), Journal of Molecular Biology 2000;299:1113–1119.

[69]. Glasscock JM, Zhu YJ, Chowdhury P et al. Using an amino acid fluorescence resonance energy transfer pair to probe protein unfolding: Application to the villin headpiece subdomain and the LysM domain, Biochemistry 2008;47:11070–11076.

[70]. Cossio P, Marinelli F, Laio A et al. Optimizing the Performance of Bias-Exchange Metadynamics: Folding a 48-Residue LysM Domain Using a Coarse-Grained Model, Journal of Physical Chemistry B 2010;114:3259–3265.

[71]. Mesnage S, Dellarole M, Baxter NJ et al. Molecular basis for bacterial peptidoglycan recognition by LysM domains, Nature Communications 2014;5:4269.

[72]. Nickson AA, Stoll KE, Clarke J. Folding of a LysM domain: Entropy-enthalpy compensation in the transition state of an ideal two-state folder, Journal of Molecular Biology 2008;380:557–569.

[73]. Gao R, Stock AM. Molecular strategies for phosphorylation-mediated regulation of response regulator activity, Current Opinion in Microbiology 2010;13:160–167.

[74]. Bourret RB. Receiver domain structure and function in response regulator proteins, Current Opinion in Microbiology 2010;13:142–149.

